# Resolution of a human super-enhancer by targeted genome randomisation

**DOI:** 10.1101/2025.01.14.632548

**Authors:** Jonas Koeppel, Pierre Murat, Gareth Girling, Elin Madli Peets, Mélanie Gouley, Valentin Rebernig, Anant Maheshwari, Jacob Hepkema, Juliane Weller, Jenie Hannah Johnkingsly Jebaraj, Ronnie Crawford, Fabio Giuseppe Liberante, Leopold Parts

**Author notes:** Contributed equally (Jonas Koeppel, Pierre Murat).

## Abstract

Human gene expression is controlled from distance via enhancers, which can form longer ‘super-enhancer’-regions of intense regulatory activity. Whether super-enhancers constitute a separate regulatory paradigm remains unclear, largely due to the difficulty of dissecting the contributions and interactions of individual elements within their natural chromosomal context. To address this challenge, we developed enhancer scrambling, a high-throughput strategy to generate stochastic inversions and deletions of targeted enhancer regions by combining CRISPR prime editing insertion of symmetrical loxP sites with Cre recombinase-induced rearrangements. We applied our approach to dissect a distal super-enhancer of the *OTX2* gene, generating up to 134 alternative regulatory configurations in a single experiment, and establishing how they drive gene expression and chromatin accessibility, as well as the individual contributions of its elements to this activity. Surprisingly, the presence of the sequence containing a single DNase I hypersensitive site predominantly controls *OTX2* expression. Our findings highlight that enhancer-driven regulation of some highly expressed, cell-type-specific genes can rely on an individual element within a cluster of non-interacting, dispensable components, and suggest a simple functional core to a subset of super-enhancers. The targeted randomisation method to scramble enhancers can scale to resolve many super-enhancers and human gene regulatory landscapes.

## Introduction

For almost every human gene, several distal DNA-encoded regulatory elements control its transcription in time and space. Mutations disrupting enhancers can dysregulate gene expression, and lead to disease^1,2,3,4^. Large clusters of enhancers that drive the expression of genes which define cell identity have been termed ‘super-enhancers’^5–7^, and their destabilisation can impact development^5^ and tumorigenesis^8,9,10^. A fundamental outstanding question in gene regulation, particularly for super-enhancers, is how a regulatory architecture functions as a combination of its constituent parts and across distance^8,11,12^. Answering this question requires extensive data on enhancer affinity for transcription factors, as well as quantitative estimates of how the number, arrangement, spacing, and orientation of enhancers influence the expression of associated genes.

Current technologies limit the complexity of regulatory landscapes that can be explored at scale. Regulatory potential of individual sequences can be assessed through transcription factor binding experiments^13,14^ and massively parallel reporter assays^15,16^, however, these methods often fail to preserve the native genomic context. Genomic manipulation methods retain this context but are typically confined to single specific modifications, such as deletions, inactivations, or activations^17,18^. Both approaches usually target single elements^19,20^, with only limited attempts to examine effects of paired elements or spacing variations. To investigate how factors like combinatorial arrangement, orientation, spacing, and interactions between elements influence gene expression, synthetic reconstruction of large gene loci has been instrumental^21,22,23,24,25^. However, this process is complex, challenging to scale, and can require specialised infrastructure across various model organisms, from yeast to mammalian cells. Though a few exemplar cases have been thoroughly studied, functional data on human super-enhancers remains scarce, with comprehensive information available for only two clusters regulating the *α-globin* and *Wap* loci^26,27,28^_._

A convergence of new technologies now enables large-scale generation of data on regulatory configurations within native chromatin contexts^29^. For example, synthetic DNA transposon ‘hopping’ has been applied to relocate functional elements within the genome and study their effects when placed in new genomic sites^30,31^. CRISPR-based methods, notably CRISPR prime editing, expand these capabilities by enabling precise insertion of short sequences, such as loxPsym sites, at specific genomic locations^32,33^. These inserted sites can then serve as substrates for Cre recombinase to create new genomic configurations through stochastic deletion or inversion of intervening DNA in both yeast^34^ and human cells^35^. Advances in long-read sequencing further enhance this approach, providing detailed long-range mapping of these rearrangements^36^ while also revealing CpG methylation patterns without PCR-induced biases^37,38^. Additionally, accessible chromatin regions can be marked with N^6^-Methyladenosine (m6A), quantified from nanopore sequencing signals, and linked to the generated DNA variants^39^.

Here, we leverage these technologies to scramble human regulatory regions, and generate variation into regulatory architectures at scale. Our focus is a ∼75 kb domain upstream of the *OTX2* gene, which contains multiple enhancer candidates within a super-enhancer. We combine prime editing with Cre-lox programmed recombination to construct synthetic genomic architectures and assess their effects on gene expression in parallel using long read single molecule sequencing. With a small set of experiments yielding hundreds of distinct regulatory configurations, we examine the super-enhancer contribution to *OTX2* expression in HAP1 cells, quantify the roles of its individual elements, and map the corresponding changes in locus accessibility and methylation. Notably, we find that a single DNase I hypersensitive site predominantly regulates *OTX2* expression. Our work provides both a framework for high-throughput enhancer function analysis and new insights into super-enhancer biology.

## Results

We devised enhancer scrambling, a targeted randomisation strategy that generates multiple alternative gene regulatory architectures within a single experiment, enabling comparison of their gene expression potential (**Fig. 1a**). This method uses prime editing to insert loxPsym sequences between regulatory elements of a fluorescently tagged target gene. Upon delivering Cre recombinase, these sequences undergo recombination, resulting in stochastic deletions, inversions, or combinations of both, generating a diverse cell pool with a random regulatory landscape in each cell.

**Figure 1.**
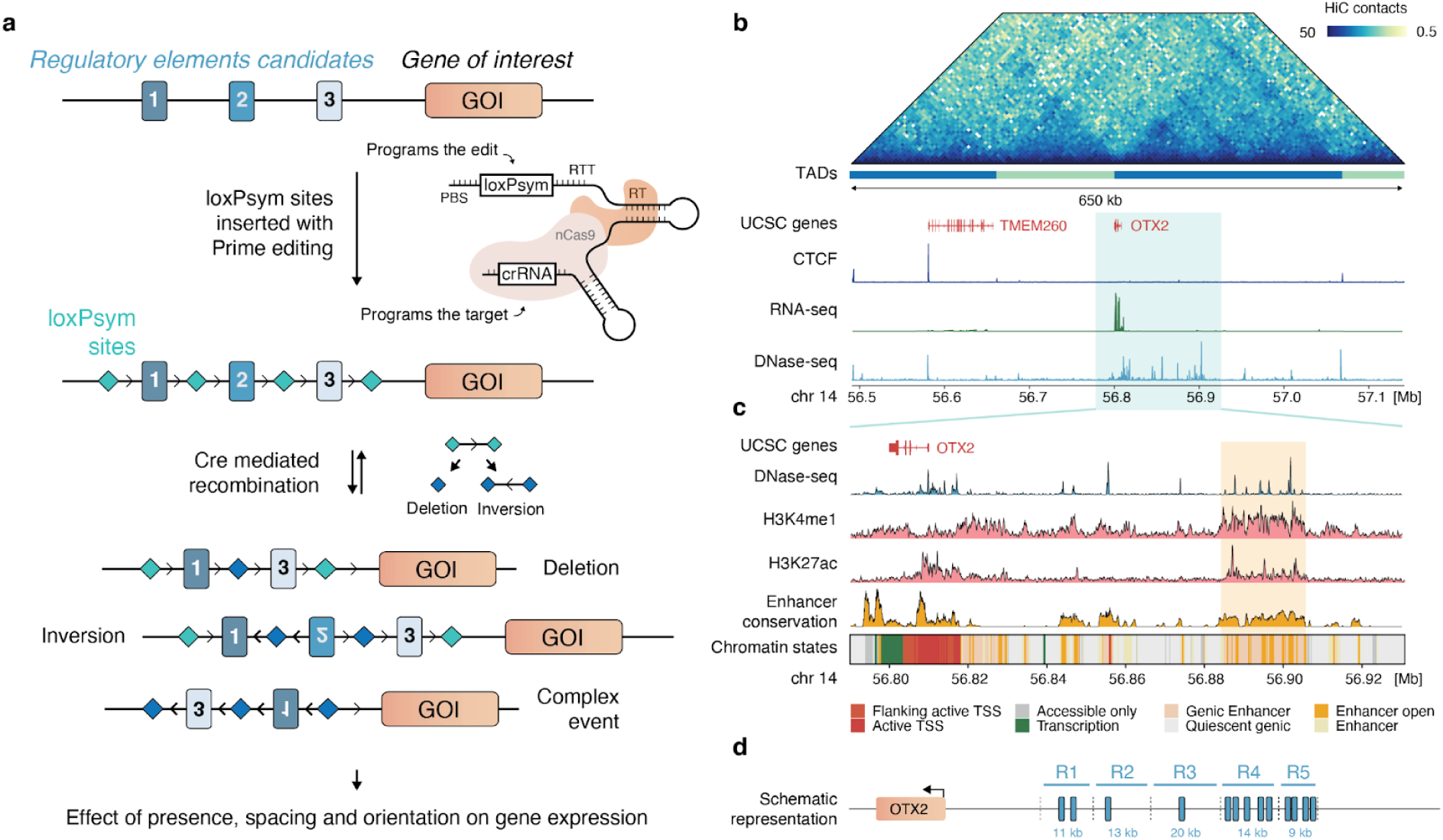
A prime editing-based randomisation approach for dissecting architecture of *OTX2* regulation. **(a)** Overview of enhancer scrambling. **(b)** *OTX2* is the only gene expressed within its topologically associated domain (TAD) in HAP1 cells. A ∼75 kb region (x-axis) upstream of *OTX2* has frequent DNase hypersensitivity (DHS) peaks (bottom track) and increased interactions to the gene (top heatmap). **(c)** Genomic tracks (y-axis) of DNase-seq (top), histone marks (H3K4me1 and H3K27ac; middle), conservation, and ChromHMM segmentation (bottom, colors) with a potential distal super-enhancer highlighted (orange box). **(d)** Schematic representation of the *OTX2 cis*-regulatory region and the enhancers within (blue rectangles), illustrating the design for loxPsym site insertion (dashed lines) that generates regulatory domains R1-R5.

We applied enhancer scrambling to investigate the *cis*-regulation of the *OTX2* gene. *OTX2*, a homeobox transcription factor, is highly expressed but non-essential in HAP1 cells (**Supplementary** Fig. 1a,b), with expression levels varying across different cell lines and tissues^40^. It is the only expressed gene within a megabase region on chromosome 14 in HAP1 cells and the only gene within its topologically associating domain (**Fig. 1b**). These features suggest the presence of distal regulatory mechanisms and reassure that structural variants within this locus would not affect cellular fitness or the expression of neighbouring genes. We identified a ∼75 kb region upstream of *OTX2* containing 13 DNase I hypersensitive peaks, along with H3K4me1 and H3K27ac marks, hallmarks of distal enhancer activity (**Fig. 1c**). A subset of these peaks forms a ∼22 kb cluster, which qualifies as a super-enhancer, distinguished by closely spaced genomic regions exhibiting enhancer signatures, including H3K27ac (**Supplementary** Fig. 1c)^6^. Using enhancer conservation data across 127 cell types from the Roadmap Epigenomics Consortium, we defined five domains of potential regulatory activity (“R1”-”R5”, **Fig. 1d** and **Supplementary** Fig. 1d), that we then set to randomise to explore their impact and interactions.

To rearrange the regulatory region, we first tagged the *OTX2* gene in HAP1 cells with the mScarlet fluorescent marker and used CRISPR prime editing, followed by two rounds of subcloning, to generate clonal cell lines with three or six loxPsym sites inserted at the R1-R5 domain boundaries (‘OTX2-loxp3’ and ‘OTX2-loxp6’ lines; **Fig. 2a** and **Supplementary** Fig. 2a). We then induced recombination in the clonal cell lines by delivering Cre recombinase (**Methods**) and allowed *OTX2* expression to stabilise over the course of one week. In a subset of treated cells, OTX2-mScarlet abundance decreased (**Fig. 2b** and **Supplementary** Fig. 2b), indicating the formation of new regulatory architectures that lead to reduced *OTX2* expression. We derived two single clones from the population of scrambled OTX2-loxp3 cells, established that they have deletions of both the R4 and R5 regions, and measured a 52% reduction in their *OTX2* transcripts and no changes in neighbouring gene expression by RNA sequencing (**Supplementary** Fig. 2c), confirming the causal role of the super-enhancer.

**Figure 2.**
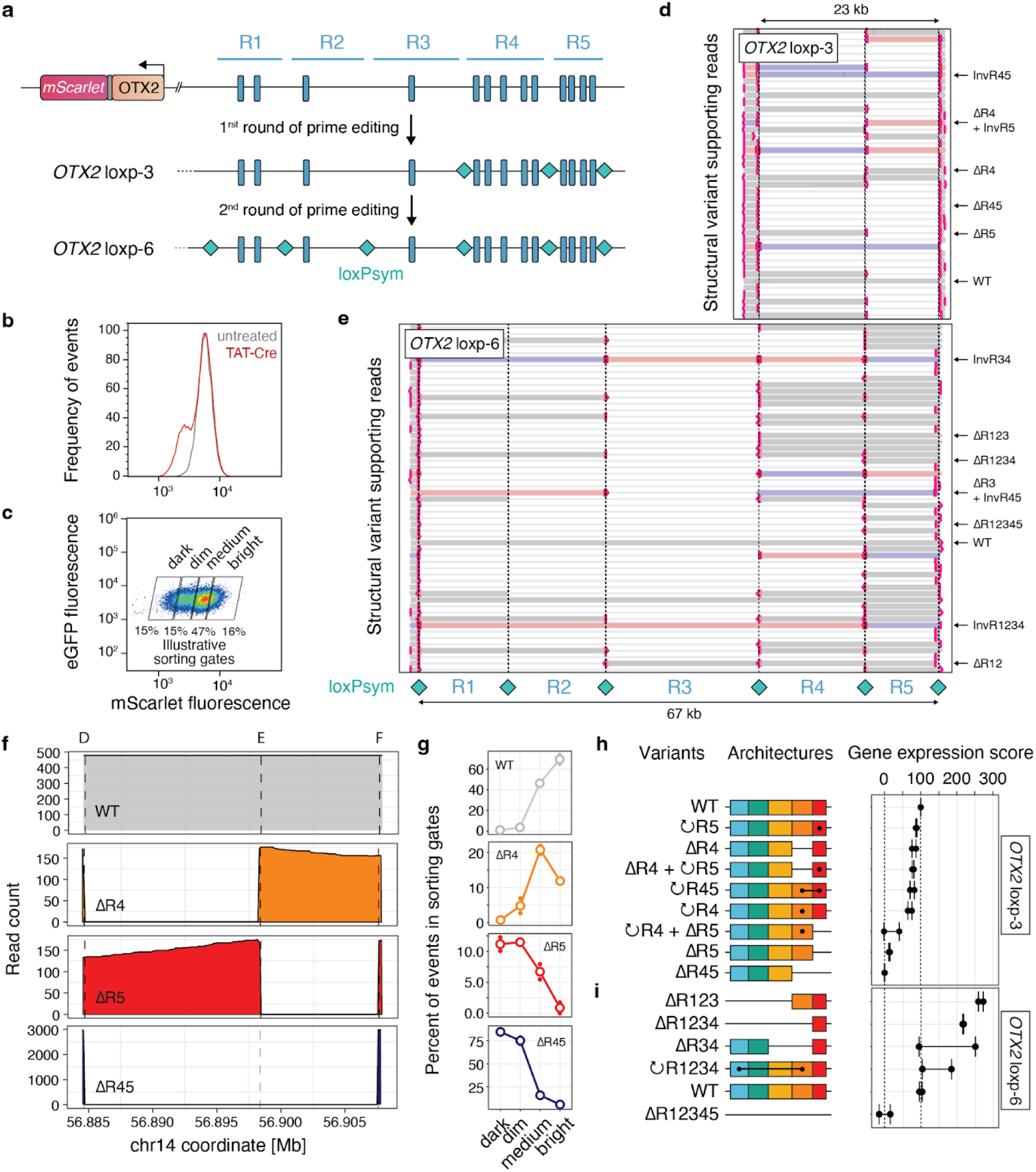
Cre induction randomises the *OTX2* regulatory architecture, facilitating functional analysis of enhancer combinations. **(a)** Schematic of the prime editing strategy used to generate clonal HAP1 cell lines OTX2-loxp3 and OTX2-loxp6, containing three and six loxPsym sites (teal diamonds) respectively, that separate enhancers (blue rectangles). **(b)** Frequency (y-axis) of mScarlet fluorescence (x-axis) in OTX2-loxp3 cells with (red) and without (grey) Cre recombinase treatment. **(c)** Example of sorting gates (boxes) used to isolate cells (markers and colours) with varying levels of OTX2-mScarlet expression (x-axis), with eGFP fluorescence (y-axis) as a control. **(d)** Cas9 enrichment combined with nanopore sequencing identifies structural variants within mixed-genotype populations derived from the OTX2-loxp3 cell line in individual reads (y-axis). Red and blue colours: different strands. Arrows and vertical lines: direction and breaks within the reads. **(e)** As **(d)**, but for the OTX2-loxp6 cell line. **(f)** Read count (y-axis) along the genome (x-axis) for architectures (panels, labels) identified from long-read sequencing. **(g)** Percentage of events (y-axis) in sorting bins with increasing levels of *OTX2* expression (x-axis) for different genotypes (panels). Markers: biological replicates. **(h)** Relative gene expression (x-axis) for different architectures (y-axis) in OTX2-loxp3 cells. Markers: biological replicates; dashed vertical lines: wild-type and full-deletion values; dashed horizontal lines: deleted sequences; dots and connecting lines: inverted sequences; colours: regulatory regions. **(i)** Same as **(h)**, but for OTX2-loxp6 cells.

We next aimed to separate the cells based on expression, and to identify the regulatory architectures generated by scrambling. To do so, we sorted the cells based on mScarlet fluorescence into dark, dim, medium, and bright categories, and confirmed the expression levels remained stable after an additional seven days in culture (**Fig. 2c**, **Supplementary** Fig. 2d,e). We then performed Cas9-enriched nanopore long-read sequencing on high molecular weight DNA extracted from the sorted cell pools (**Methods**), and identified a range of sequence variants. The rearrangements included deletions, inversions, and combinations of both, with breakpoints at the loxPsym sites in both the OTX2-loxp3 and OTX2-loxp6 cell lines (**Fig. 2d** and **2e**), spanning 23 kb and 67 kb, respectively. We called structural variants (**Methods; Supplementary** Fig. 3) and grouped reads to identify 7 and 6 distinct synthetic architectures generated in the OTX2-loxp3 and OTX2-loxp6 experiments (**Supplementary** Fig. 4) with consistent read coverage (**Fig. 2f**) that supported calculating variant frequencies in each bin (**Fig. 2g**).

Having genotyped the reads, we then quantified the impact of each synthetic architecture on gene expression. To facilitate quantitative comparisons, we calculated a weighted average of the genotype frequency across expression bins, scaled it to range between 0 (equivalent to expression of the largest deletion variant) and 100 (equivalent to wild type expression), and computed this score for the architectures with sufficient coverage (**Fig. 2h** and **2i, Methods**). In OTX2-loxp3 cells, the unedited wild-type configuration was more frequent in the brighter bins associated with higher gene expression levels, while deletions of the R5 domain led to a decrease in *OTX2* expression (**Fig. 2g** and **Supplementary** Fig. 5a). Deletion of only the R4 region, and inversions of single or multiple domains had minimal impact (**Fig. 2i**). Likewise, variants combining inversions and deletions showed expression levels similar to those with deletions alone. For example, cells with both an R4 deletion and an inversion of the R5 domain (ΔR4 + ↻R5) exhibited expression patterns comparable to those with only the R4 deletion (ΔR4). Interestingly, deletions of upstream domains (R1, R2, and R3), introduced by scrambling in the OTX2-loxp6 cell line, led to an increase in gene expression compared to the wild-type configuration (**Fig. 2i** and **Supplementary** Fig. 5b). Taken together, deleting R5 leads to lower *OTX2* expression, while bringing it closer to the gene enhances its expression, establishing this domain of the super-enhancer as a key *OTX2* regulator.

Long-read sequencing of unamplified genomic DNA enables assessment of scrambling effects on DNA accessibility by treating chromatinised DNA with a non-specific adenine methyltransferase (**Methods**)^39^. We generated DNA accessibility profiles from WT reads of OTX2-loxp3 and OTX2-loxp6 cells, and found good agreement with publicly available DNase-seq accessibility data from unedited HAP1 cells (**Supplementary** Fig. 6a)^41^. Since WT reads were not systematically more accessible in the bright than the dark sorting gate overall (median adenine methylation of 1.63% vs 1.67%, **Supplementary** Figure 6b,c), or between any other two gates (**Supplementary** Figure 6d), we combined reads across all sorting gates to compare the accessibilities between synthetic architectures. We then quantified DNA accessibility at the five DHS peaks within the *OTX2* super-enhancer. Deletions that decreased or increased expression had reduced or elevated accessibility at nearby sites respectively (**Fig. 3c**). For instance, deletion of the R5 domain reduced median accessibility at DHS peaks in the adjacent R4 domain by 41%, while deletion of the accessory R1-R2-R3 region led to a 15% increase. Similarly, deletions of the R4, R1-R3, or R1-R4 regions which all move R5 closer to the transcription start site, increased accessibility at DHS peaks in the R5 domain by a median of 82-100% (**Fig. 3d**).

**Figure 3.**
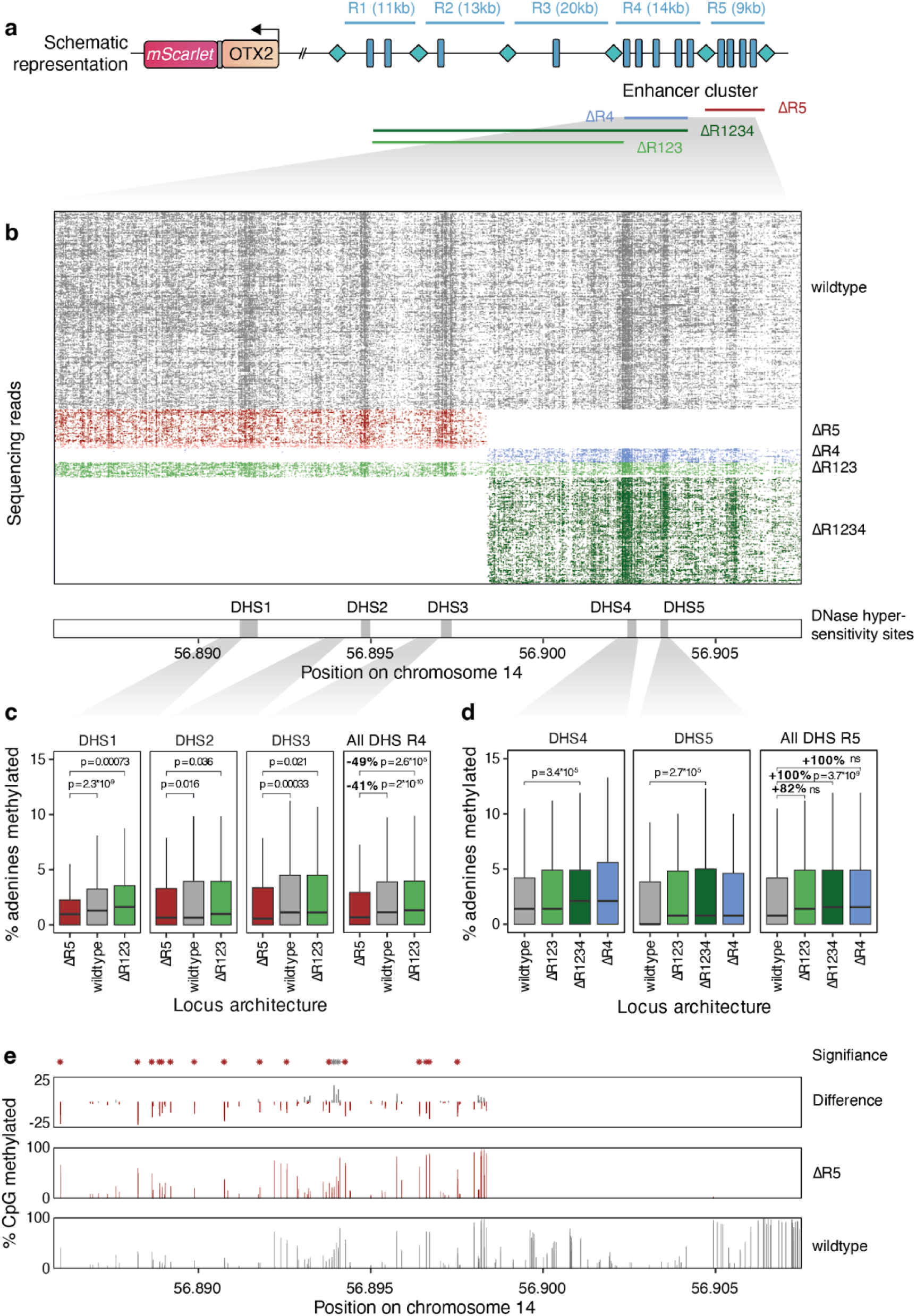
The *OTX2* super-enhancer regulates DNA accessibility. **(a)** Schematic of the deletion variants analyzed to assess the role of the R4:5 super-enhancer in DNA accessibility. **(b)** Individual nanopore sequencing reads (rows) with m6A-modified bases (dashes) for the WT (grey) and deletion variant (colors) architectures. Large white regions: missing due to deletions. **(c)** Percentage of m6A modifications (y-axis), used as an indicator of DNA accessibility, at individual DHS peaks of the R4 domain (left three panels), as well as combined (right panel) for different locus architectures. Box plots: medians and interquartile ranges. *P*-values: two-tailed Student’s t-test. **(d)** As **(c)** but for R5 domain DHS peaks. **(e)** The percentage of CpG island methylation across the R4 and R5 domains for wild type (bottom panel, grey), the ΔR5 configuration (middle panel, red), and their difference (top panel). Red stars: positions of significant increase in ΔR5 (absolute difference of > 10%, Benjamini–Hochberg corrected p < 0.01; two-tailed Student’s t-test); grey stars: positions of significant decrease.

To corroborate these findings, we analyzed CpG methylation from the same reads. The R5 domain deletion led to increased CpG methylation across the remaining CpGs in the enhancer cluster (27/29 CpGs with significant differences to the WT were more methylated when the R5 domain was deleted), indicative of less active chromatin (**Fig. 3e**). Together, these results suggest that the R5 domain presence promotes DNA accessibility and chromatin activity, so that deletions within the super-enhancer domains induce changes in average DNA accessibility as well as CpG methylation.

Having established the functional role of the *OTX2* super-enhancer, we next addressed three limitations of our initial approach: (i) generating cell lines with multiple loxPsym integrations took months for editing, subcloning, and genotyping; (ii) recombination reactions in clonal cell lines frequently resulted in complete locus deletions; and (iii) Cas9 enrichment sequencing produced limited number of reads. To expedite enhancer scrambling and increase variant diversity, we co-transfected a pool of epegRNAs from the OTX2-loxp6 design (**Fig. 2a**) without clonal selection, achieving an editing rate of 11 to 36% (mean 25%) across the six target sites. This approach yielded a broader range of synthetic architectures (18 architectures compared to the 14 previously identified, **Supplementary** Fig. 7) with reduced bias toward full deletions (3% of events compared to 23% before; **Supplementary** Fig. 8), and confirmed that deletion of the domain upstream of the super-enhancer leads to an increase in *OTX2* expression. To improve variant detection sensitivity, we adapted a targeted nanopore sequencing protocol using long-range PCR^36^. PCR sequencing of the 23.3 kb targeted region increased variant coverage by ∼100-fold on average, while maintaining the enrichment of variants across sorting gates and their gene expression scores (**Supplementary** Fig. 9). It is noteworthy that all nine possible synthetic architectures derived from the OTX2-loxp3 design could be characterized in this way. Overall, this optimised protocol produced synthetic architectures within six weeks from transfection to sequencing, enhanced variant diversity, and improved detection sensitivity.

We next applied our optimized protocol to fine-map the elements of the R4-R5 super-enhancer region. By transfecting epegRNAs into OTX2-loxp3 cells, we introduced an additional loxPsym site within the R4 region and three within the R5 region (“OTX2-loxp7” design, **Fig. 4a**, **Methods**), achieving editing rates of 5% to 36% after two transfection rounds (**Supplementary** Fig. 10a). Following scrambling (**Supplementary** Fig. 10b), fluorescence sorting (**Supplementary** Fig. 10c), long-range PCR, and nanopore sequencing, we observed a variety of amplicon lengths (**Fig. 4b** and **4c**), and confirmed a ∼50% reduction in *OTX2* expression in the darkest sorting gate (**Supplementary** Fig. 10d). Using an optimized structural variant caller (**Methods**), we identified 134 distinct architectures (**Supplementary** Fig. 11), including the WT configuration, 19 out of the 21 possible simple deletions, and an additional 114 combinations involving simple inversions, multi-breakpoint deletions, and deletions combined with inversion events. We also uncovered complex rearrangements where individual elements are duplicated, resulting from recombination of the haploid genome during the S-phase of the cell cycle, or where the order of enhancer elements is altered (**Fig. 4d**). The distribution of reads across different *OTX2* expression bins revealed the contributions of individual enhancer elements. Notably, progressively larger deletions from the 5’ end of the super-enhancer significantly affected *OTX2* expression only when the R5-2 domain was disrupted (**Fig. 4e**), suggesting a critical role for this element in the function of the super-enhancer. Overall, the optimized enhancer scrambling can efficiently generate over a hundred synthetic regulatory architectures in a single experiment.

**Figure 4.**
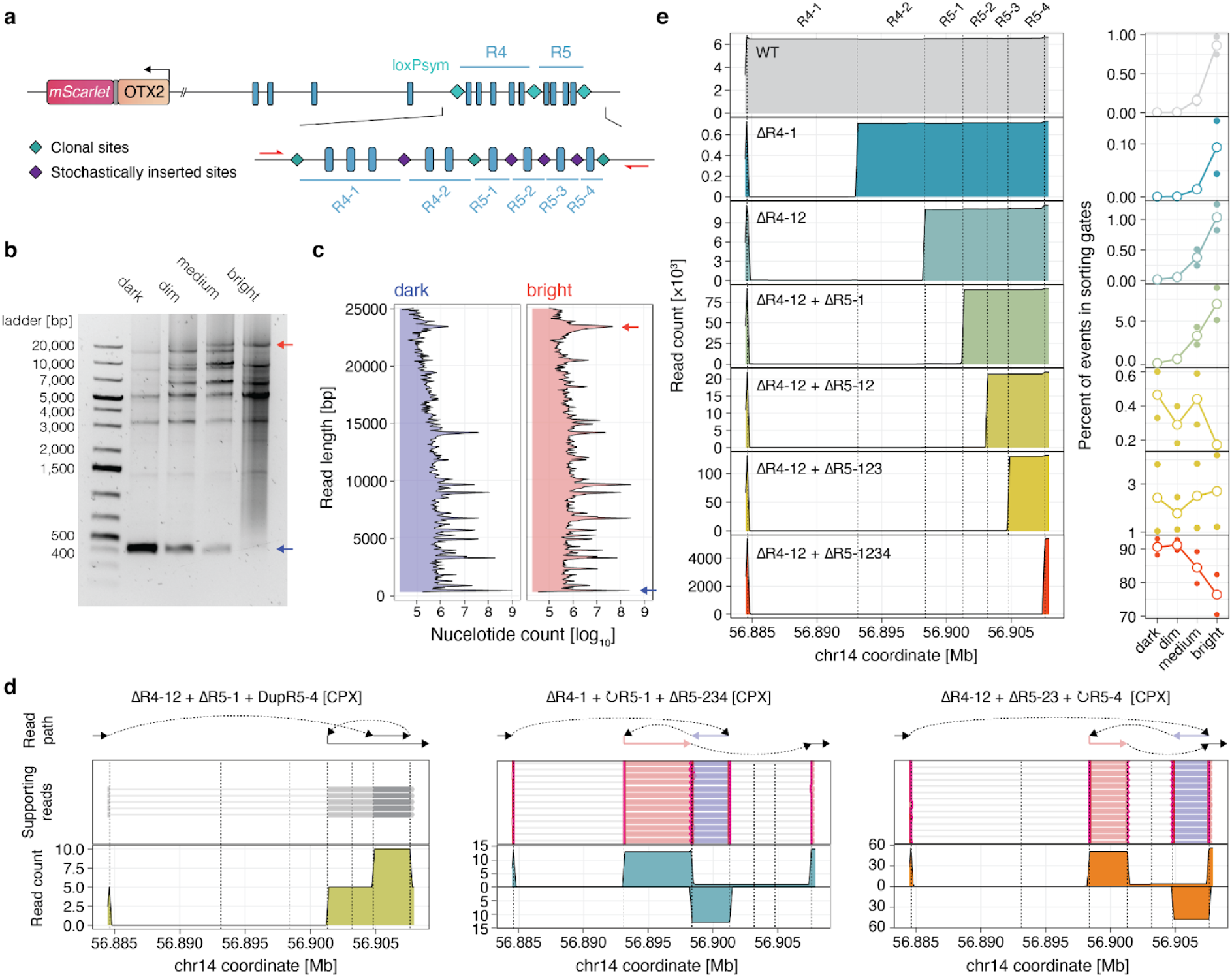
Functional dissection of the OTX2 super-enhancer activity. **(a)** Schematic of the pooled transfection approach combined with long-range PCR and nanopore sequencing to analyse the activity and synergy of short enhancer regions. **(b)** Gel electrophoresis of long-range PCR amplicons post-randomisation and sorting reveals numerous structural variants associated with *OTX2* gene expression. **(c)** Read length (y-axis) frequency (x-axis, log10-scale) in dark (left panel) and bright (right panel) gates shows discrete populations of structural variants. Red arrow: wild type sequence. Blue arrow: deletion from first to last loxPsym site. **(d)** Example architectures (out of 134) from scrambled OTX2-loxp7 cells. Read coverage (y-axis) across the *OTX2* super-enhancer region (x-axis), on the forward strand (positive values) and bottom strand (negative values) for duplication (left panel), and complex rearrangements (middle and right panel). Arrows and vertical lines: direction and breaks within the reads on the read path (top). Middle lines: individual supporting long reads (as in Fig. 2d). Dashed lines: locations of the three clonal loxPsym sites. **(e)** The impact of increasingly long deletions (panels) of the *OTX2* super-enhancer. Left: Read coverage (y-axis) across the region (x-axis). Dashed lines: locations of the three clonal and four subclonal loxPsym sites. Right: Percentage of reads (y-axis) carrying each deletion within the sorting gates (x-axis). Markers: replicate values.

Altogether, we could quantify the gene expression impact of 61 of the 134 synthetic architectures from the scrambled OTX2-loxp7 cells (**Fig. 5a**, **Supplementary** Fig. 12, **Methods**), with several deletions substantially reducing *OTX2* expression. Inversions had minimal effects, with a strong correlation between the gene expression scores of architectures with inversions and those of the corresponding architectures without inversions (Pearson’s R=0.87, *P*=4.2x10^-11^, **Fig. 5b**). These observations suggest that deletions of functional elements, rather than changes in the distances between the super-enhancer elements and the gene promoter, impact gene expression in this setting. This effect is likely due to the 80kb distance between the super-enhancer and gene promoter, compared to the spans of the domain inversions within the 26kb locus.

**Figure 5.**
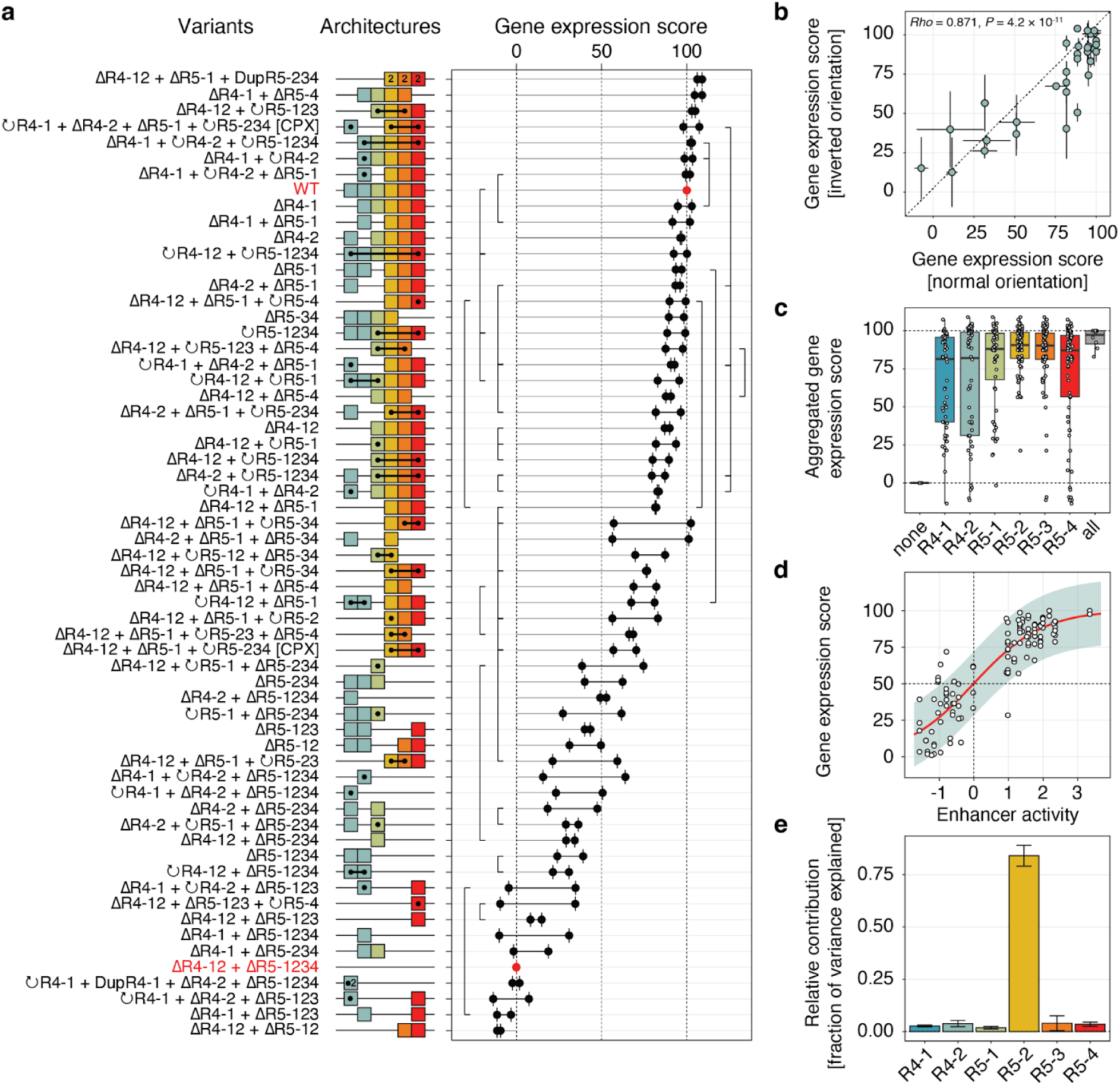
Determinants of *OTX2* super-enhancer activity. **(a)** Gene expression (x-axis) for all 61 high-confidence regulatory architectures (y-axis) from scrambled OTX2-loxp7 cells. Labels: annotation of deletions (Δ) and inversions (↻) of the regions, e.g. full deletion: ΔR4-12 + ΔR5-1234. CPX: label for rearrangements that no longer follow the original order of the domains. Red text: labels for full deletion and wild type architectures used for scaling expression values between 0 and 100 (outer dashed lines). Colored boxes: enhancer domains (R4-1, R4-2, R5-1, R5-2, R5-3, and R5-4). Black dots and connecting lines on boxes: inversions; numbers on boxes: duplications. Black dots (right): replicates. Vertical connecting lines (right): link two architectures that differ by a single inversion. **(b)** Replicate average gene expression score of inversion pairs from (a) (markers), where the inverted sequence is in normal orientation (x-axis) or inverted orientation (y-axis). Whiskers - replicate range. **(c)** Gene expression scores (y-axis) from the 61 architectures in (a), stratified by presence of each of the regulatory domains (x-axis). Box: median and quartiles; markers: individual values. **(d)** A linear-logistic regression model fit (red line) of gene expression (y-axis) from domain presence captured as enhancer activity (x-axis, **Methods**). Coloured area: 95% confidence interval; white points: individual values for each architecture. **(e)** Fraction of variance explained (y-axis) by each enhancer domain (x-axis) in the model from (d). Whiskers: replicate range.

Architectures retaining the R5-2 and R5-3 domains had consistently high gene expression, whereas architectures with other domains showed more variable outcomes (**Fig. 5c**). To quantify the contribution of individual regions, we tested a range of generalised linear models and selected a linear-logistic model as the best fit (**Supplementary** Fig. 13, **Methods**). The selection of this model suggests that the super-enhancer operates as a two-state system, transitioning from a quiescent state to an active state upon the activation of key individual enhancer components (**Fig. 5d**). Surprisingly, the R5-2 region, which features a single DNase hypersensitive peak, accounted for ∼80% of the variance in the observed gene expression levels (**Fig. 5e**), while other regions made considerably smaller contributions. Incorporating pairwise, but not higher order interactions between regions only slightly improved the fit (**Supplementary** Fig. 13e and 13f). These findings suggest that the *OTX2* super-enhancer likely depends on a single enhancer element within the R5-2 region, with minimal interaction between its components.

Systematic screening of enhancer activity has generated large-scale data that have informed predictive models of enhancer-gene regulatory interactions. We tested whether the regulatory role of the *OTX2* super-enhancer, particularly the R5-2 region, could be predicted and quantified using such models (**Supplementary** Fig. 14a). First, we applied the Activity-by-Contact (ABC) model,^42^ which integrates chromatin state data with 3D physical interactions to assess enhancer-gene relationships, to HAP1 cells. Despite HiC evidence suggesting a connection (**Fig. 1b**), ABC failed to detect the interaction between the *OTX2* super-enhancer and its target gene. We then employed the ENCODE-rE2G model^42,43^, a supervised classifier that builds on ABC, which did identify the R5-2 to *OTX2* interaction, albeit with low confidence. Next, we assessed whether the variable gene expression effects of synthetic landscapes could be predicted using Borzoi^44^, a state of the art AI model for gene expression (**Methods**, **Supplementary** Fig. 14b). Deletion variants were predicted to have minimal impact on *OTX2* expression (expression range from 1.5 to 1.6; **Supplementary** Fig. 14c), with weak correlations to observed values in all tested cell types (maximum Pearson’s R = 0.16, average = -0.16; **Supplementary** Fig. 14d). These results suggest that while some models can predict distal regulatory interactions with low confidence, additional data and refined models are necessary for more accurate predictions of their contribution.

Finally, we aimed to characterise the mechanism by which the R5-2 domain of the super-enhancer regulates *OTX2* expression. We identified five transcription factors expressed in HAP1 cells (FOXA1, FOXK2, RFX5, YY1, and MYB) with 12 predicted binding sites specifically in the region using maxATAC^45^ (**Methods**, **Supplementary** Fig. 15a,b). We designed epegRNAs to induce 6-8nt small deletions at the sites (**Fig. 6a, Methods**) and pool transfected them, resulting in a population of cells with reduced fluorescence (**Supplementary** Fig. 15c), suggesting that these sites contribute to *OTX2* expression. We then enriched these cell populations using FACS (**Fig. 6b**), and observed the expected short deletions (**Supplementary** Fig. 15d) alongside longer deletions spanning the entire amplicons (**Fig. 6c**), attributed to the concurrent prime editing activity of the 12 epegRNAs. The deletions at 5/12 predicted sites, spanning ∼150 bp of the R5-2 ATAC-seq peak, were significantly associated with *OTX2* expression (**Supplementary** Fig. 15d and **Fig. 6d**). This short DNA sequence contains four of the six predicted FOXA1/FOXK2 binding sites, indicating a potential role for these transcription factors in regulating OTX2 expression. Taken together, these findings underscore the functional significance of the single accessible site of the R5-2 domain and highlight short 6–8 nt sequences overlapping predicted transcription factor binding sites as regulators of *OTX2* expression.

**Figure 6.**
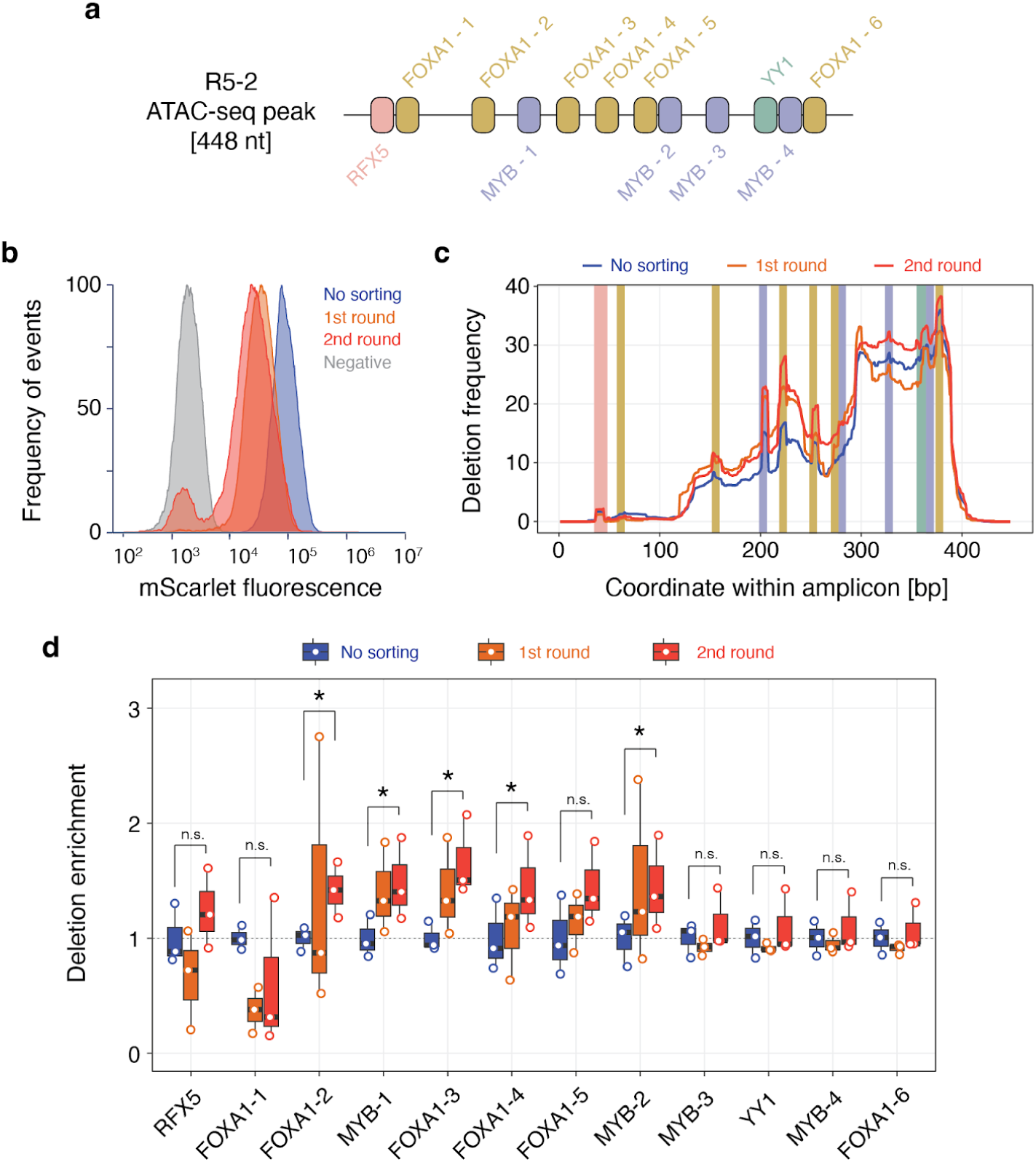
Enhancer scramble reveals distal transcription factor binding sites associated with *OTX2* expression. **(a)** A single ATAC-seq peak within the R5-2 domain, identified as the central active element of the *OTX2* super-enhancer, contains consensus DNA binding sequences (boxes) for the transcription factors RFX5, FOXA1/FOXK2, MYB, and YY1 (colors). Predicted binding sites for FOXK2 overlaps with those identified for FOXA1. **(b)** Frequency of cells (y-axis) with different mScarlet expression (x-axis) after transfection with 12 epegRNAs, before sorting (blue), after one round of sorting (pink), and after two rounds of sorting (red), compared to mScarlet-negative cells (grey). **(c)** Deletion frequency (percent, y-axis) along the R5-2 ATAC-seq peak (x-axis) for amplicons generated from unsorted (blue) and sorted cells (pink, red). Coloured boxes: targeted transcription factor binding sites; colors as in **(a)**. **(d)** Enrichment of deletions (y-axis) at targeted transcription factor binding sites (x-axis) compared to unsorted cells for, different numbers of sorting rounds (bars, colors). Colors as in (b). *P*-values: two-tailed Student’s t-tests; n.s.: not significant, \**P* < 0.05.

## Discussion

We presented enhancer scrambling, a novel approach to investigate gene regulatory regions in their native context by combining multiplexed prime editing with Cre-lox technology to induce stochastic rearrangements. This strategy enabled the generation of hundreds of unique regulatory architectures in just a few experiments, elucidating the *cis*-regulation of *OTX2* by non-coding elements within a super-enhancer. We demonstrated that this super-enhancer contributes ∼50% to *OTX2* expression in a distance-dependent manner. Single-molecule long-read sequencing revealed how enhancer deletions impact chromatin accessibility and CpG methylation across the locus. By optimizing our design, we generated 134 and analyzed 61 synthetic super-enhancer variants in a single experiment within weeks, bypassing the need for clonal selection of engineered cells, and identified a key ∼150bp sequence within a single accessible chromatin region by mutating its transcription factor binding sites. Together, our approach streamlines the study of gene expression regulation and elucidates the regulatory mechanisms of a model super-enhancer.

The expression of *OTX2* relied primarily on a single enhancer element, with its position and orientation within the wider super-enhancer having limited effect. This simplicity is surprising, given current models of super-enhancer biology, which suggest that individual enhancers can act additively, redundantly, or synergistically, depending on the context and cell type^26,27,46^. In these studies and subsequent analyses of related data^11^, super-enhancer activity was best explained by a two-state model, with contributions from each element. However, limited data precluded rigorous testing of interaction effects between individual elements. With our larger dataset, we found minimal contributions beyond the independent effects of each element, suggesting that the *OTX2* super-enhancer functions as a cluster of independent elements in HAP1 cells, with largest contribution from the R5-2 domain. While we focused on the *OTX2* super-enhancer, our optimised protocol can be applied to investigate other enhancer clusters to test the generality of this observation.

Super-enhancers and their target genes respond to cell state transitions, both during development and in reaction to transcriptional or environmental stimuli^47,48^. Interactions between individual enhancer components may thus become active during the temporal formation of the *OTX2* super-enhancer or in response to specific external conditions. Our framework is well-suited to investigate these dynamic regulatory relationships. Extending this approach across diverse cell types and conditions will, however, require further optimisation of prime editing, which remains challenging in key cell systems such as pluripotent stem cells^49,50^. Iterations of scrambling, adding further loxPsym sites to regions with high signal, increase the number of structural variants to investigate complex locus control regions at higher resolution, as we demonstrated by further dissecting the R4 and R5 regions.

Enhancer scrambling captures regulatory complexity more effectively than pooled DNA oligonucleotide-based assays like MPRAs^14^ and is more scalable than resynthesizing multiple regulatory landscape versions^22^. Nonetheless, some considerations for scaling remain. The repertoire of generated architectures is biased by technical covariates, with the read count for each architecture correlated with the number of recombination events required for their generation (**Supplementary** Fig. 16a), the length of the structural variant (**Supplementary** Fig. 16b), and the frequency of loxPsym insertion events that produced these variants (**Supplementary** Fig. 16c). Downsampling experiments suggest that some of these biases can be overcome by sufficient sequencing (**Supplementary** Fig. 16d), but future experimental and computational work will address how judged control of the biases can steer variant generation towards desired distributions^51^.

Saturation genome engineering in HAP1 cells is setting new standards for measuring clinically relevant effects of coding variants^52–56^, and we propose that it holds equal promise for uncovering key principles of gene regulation and the impact of non-coding mutations. Although AI-powered computational models provide critical insights into enhancer activity and function, they could not yet capture the impact of mutations we generated, and still require experimental validation to fully capture the intricate effects of enhancer presence, spacing, and orientation on gene expression. Our method provides a robust, scalable framework for addressing these challenges by systematically exploring a vast range of regulatory element variants within their native chromatin context. We anticipate that enhancer scrambling will help uncover fundamental principles of gene expression by exploring a previously unreachable diversity of regulatory element configurations.

## Author contributions

Designed experiments: JK, PM, JW, RC, FGL, MG, LP. Performed experiments: JK, PM, EMP, MG, VR, GG, JW, JHJJ. Analyzed data: JK, PM, AM, JH, MG, VR. Wrote paper: PM, JK, LP with input from all authors. Conceived project: LP. Supervised project: JK, PM, LP.

## Acknowledgements

All authors were supported by Wellcome (220540/Z/20/A). We thank Johan Henriksson for suggesting the pooled screening approach, and Ben Lehner and Jussi Taipale for comments on the text. We acknowledge the laboratories of John Stamatoyannopoulos, Michael Snyder, Erez Aiden, and Jesse Engreitz involved in the generation of the ENCODE datasets used.

## Materials and Methods

### Cell culture

HAP1 *ΔMLH1* cells expressing a doxycycline-inducible prime editor^11,33^ were cultured in IMDM (Invitrogen) with 10% FBS (Invitrogen) and glutamine and penicillin-streptomycin (Invitrogen) at 37 °C and 5% CO2.

### HAP1 expression data in cancer cell lines

Cancer cell line gene expression data were downloaded from the Cancer Cell Line Encyclopedia Data portal (https://sites.broadinstitute.org/ccle/datasets; 22Q2 release^57^).

### Enhancer and super-enhancer definition

All ENCODE^41^ datasets were downloaded from the ENCODE portal^41,58^. Enhancers were defined as DNase I hypersensitive sites (ENCODE accession: ENCSR895KTN) with positive ChIP-seq signals for H3K4me1 (ENCODE accession: ENCSR450JTP) and H3K27ac (ENCODE accession: ENCSR131DVD) in HAP1 cells. As previously described^6^, to capture dense clusters of enhancers, we allowed enhancer regions within 12.5 kb of one another to be ‘stitched’ together. We then ranked all enhancers based on increasing total background-subtracted ChIP-seq occupancy of H3K27ac and plotted the total background-subtracted ChIP-seq occupancy of H3K27ac in units of total rpm/bp (reads per million per base pair). The resulting plots revealed a clear threshold (rank = 6000 out of 6273 clusters, 2,500 rpm/bp) in the distribution of enhancers where the occupancy signal began to rise rapidly. We define enhancers above this threshold as super-enhancers, and those below it as typical enhancers, . To assess potential interactions between enhancers and *OTX2*, we consulted available HiC maps and associated analyses (ENCODE accession:

ENCFF898HRO). The CTCF ChIP-seq data is from study^59^ and is available at GEO (GSE180690). The enhancer conservation map from 127 reference human epigenomes, based on data generated by the Epigenomics Roadmap Project^60^, was produced from files reporting the core 15-state ChromHMM model, available at: https://egg2.wustl.edu/roadmap/web_portal/chr_state_learning.html.

### Tagging of *OTX2* with T2A-mScarlet

One day before transfection, HAP1 cells expressing doxycycline-inducible prime editor were seeded into a 24-well plate with 100,000 cells per well. 3 µl of 100 µM Alt-R CRISPR tracrRNA (Integrated DNA Technologies) were mixed with 3 µl of *OTX2*-targeting 100 µM Alt-R CRISPR crRNA (**Supplementary Table 1**, IDT) and heated at 95°C for 5 minutes and then cooled down to room temperature at 0.1°C per second. 0.7 µl of the complex was mixed with 0.5 µl SpCas9 Hifi V3 (30 pmol) and 0.3 µl sterile PBS to form the RNP. For transfection, 0.5 µl of RNP, 500 ng of DNA donor (Integrated DNA Technologies, **Supplementary Table 1**), and 2.5 µl of Cas9 plus reagent were mixed in one tube, and 1.5 µl CRISPRMAX reagent (Integrated DNA Technologies) was mixed with 25 µl of Opti-Mem in the other tube. The mixtures were combined, incubated at room temperature for 10 minutes, and added to the cells. Seven days post-transfection, cells were analyzed by flow cytometry, and mScarlet-positive cells were single-cell sorted. DNA was extracted from expanded colonies and correct mScarlet insertion was confirmed by amplicon PCR (**Supplementary Table 1**) and Sanger sequencing of the PCR product.

### pegRNA design

Suitable protospacers were identified using CHOPCHOP^61,62^ (Settings: hg38 assembly, CRISPR/Cas9, knockout). Spacers were filtered for ones that had a predicted efficiency score > 40 and no other targets in the genome with 1 or 2 mismatches. The first nucleotide of the spacer was adjusted to be a G for better U6 expression. pegRNAs were constructed by pairing spacers with the cr772 scaffold^63^ sequence gtttaagagctaagctggaaacagcatagcaagtttaaataaggctagtccgttatcaactcgaaaga gtggcaccgagtcggtgc and incorporating a 3′ extension. This extension included a 13 nt primer binding site (PBS), a 34 nt loxPsym site, and 30 nt of homology to the target. The sequences of the pegRNAs are provided in **Supplementary Table 2**.

### pegRNA cloning

Engineered pegRNAs were cloned using Golden Gate assembly, following the protocols outlined in^32,64^. Forward and reverse oligonucleotides were synthesised for the spacer, scaffold, and 3′ extensions (Integrated DNA Technologies or Merck). The engineered pegRNA acceptor plasmid (Addgene #174038) was linearised using BsaI, the oligonucleotides were annealed, and the scaffold was phosphorylated. The components were assembled via a Golden Gate reaction (utilising BsaI and T4 ligase) and transformed into XL10 Gold ultracompetent bacteria (Agilent). Plasmids were purified using a Miniprep kit (Qiagen), and successful assembly was verified through Sanger sequencing of the pegRNA.

### Inserting loxPsym sites into the *OTX2* super-enhancer (OTX2-loxp3 and OTX2-loxp6 cell line)

loxPsym site insertions were performed according to the following protocol. One day prior to transfection, 500,000 OTX2-T2A-mScarlet *ΔMLH1* HAP1 cells expressing a doxycycline-inducible prime editor were seeded into each well of a six-well plate. Prime editor expression was induced one hour before transfection by adding 1 µM doxycycline. For each well, 500 ng of each pegRNA plasmid (typically three constructs co-delivered) and 500 ng of a plasmid encoding BFP and puromycin resistance were transfected using Xfect™ Transfection Reagent (Takara Bio), following the manufacturer’s instructions. To counteract the silencing of genome-integrated prime editors during the continuous integration of loxPsym sites, 2 µg of the PE2 plasmid was also co-transfected. One day after transfection, 2 µg/ml puromycin was added to the cells. After three days, dead cells were removed along with the puromycin, and fresh media was added to allow for a two-day recovery period. Depending on experimental requirements, a second transfection round was performed, or 1–2 million cells were collected for downstream applications, such as FACS sorting, DNA preparation, or cryopreservation. The mixed populations were single-cell sorted to isolate final clones containing the expected integrations. Insertion efficiencies were estimated as described in the *Assaying Integration Efficiencies* section.

To generate both cell lines, we first performed two rounds of co-transfection with three pegRNAs targeting the last three loxPsym sites. Single-cell sorting was used to isolate a clone with two successfully integrated loxPsym sites. A subsequent round of transfection with a pegRNA targeting the third loxPsym integration was carried out using the same approach. From this population, a clone containing all three loxPsym sites was derived (OTX2-loxp3 clone). To create cells with six loxPsym insertions, the OTX2-loxp3 cells were transfected with three additional epegRNAs targeting the remaining sites and co-transfected with PE2-puromycin, while simultaneously inducing the endogenous prime editor with 1 µM doxycycline. The resulting mixed population was single-cell sorted to isolate a final clone containing all six integrations (OTX2-loxp6 clone). DNA was extracted from expanded colonies and correct loxPsym insertion was confirmed by amplicon PCR (**Supplementary Table 1**) and Sanger sequencing of the PCR product.

### epegRNA pooled-transfection and heterogeneous cell populations (OTX2-loxp6 and OTX2-loxp7 design)

Our optimized scrambling protocol leverages cell populations without clonal selection to enhance structural variant diversity, resulting in heterogeneous populations with varying editing rates at the targeted sites. The OTX2-loxp6 and OTX2-loxp7 cell populations were generated through co-transfection with a pool of epegRNAs. For the OTX2-loxp6 design, the starting point was OTX2-T2A-mScarlet ΔMLH1 HAP1 cells expressing a doxycycline-inducible prime editor. The OTX2-loxp7 population was derived from the OTX2-loxp3 clonal cell line. One day before transfection, 300,000 cells were seeded per well in 6-well plates. Prime editor expression was induced with 1 µM doxycycline one hour prior to transfection. Transfection was performed with a total of 5 µg of epegRNA using the Xfect™ Transfection Reagent (Takara Bio) following the manufacturer’s instructions. Each transfection mixture included 500 ng (10%) of a plasmid encoding BFP and puromycin resistance. Twenty-four hours after transfection, 2 µg/ml puromycin was added to the cells. After three days, dead cells were removed along with the puromycin, and fresh media was added to allow recovery for two days. A second round of transfection was then conducted. Finally, 1–2 million cells were harvested for downstream applications, including FACS sorting, DNA preparation, or cryopreservation. Insertion efficiencies were estimated as described in the *Assaying integration efficiencies* section below.

### Assaying integration efficiencies

DNA was extracted from cell pellets using the DNeasy Blood & Tissue Kit (Qiagen), following the manufacturer’s protocol with a single modification: 3 µl of RNase A (New England BioLabs) was added to a mixture of 180 µl PBS and 20 µl proteinase K during the resuspension of the cell pellets. To amplify the respective insertion sites, 10–100 ng of extracted DNA were used as input for PCR. Amplification was performed for 30 cycles using the Q5 High-Fidelity PCR Master Mix (New England BioLabs). The resulting PCR products were purified using either the QIAquick PCR Purification Kit (Qiagen) or the Monarch PCR & DNA Purification Kit (New England BioLabs) and analyzed via capillary gel electrophoresis on a DNA High Sensitivity Chip (Agilent Bioanalyzer). Insertion efficiencies were estimated by calculating the molar ratio of the higher molecular weight DNA band (representing the loxPsym insertion) to the total of the unedited and edited DNA bands. The primers used to amplify the junctions are listed in **Supplementary Table 1**.

### Scrambling OTX2-loxp3 cells and OTX2-loxp7 cell populations

The day before the experiment, 500,000 cells per well were seeded into a 6-well plate containing 1.5 ml of media. On the experiment day, 2 µM of purified TAT-Cre protein (Cambridge University Biochemistry Department) was added to the cells. They were incubated at 37 °C for 4–6 hours, after which the media was replaced, and the cells were allowed to recover overnight. The next day, the experiment was divided into two replicates, with the cells expanded over 14 days to allow gene expression to reach a steady state. Subsequently, the cells were sorted into four gates based on OTX2-mScarlet fluorescence intensity, targeting 10,000 to 100,000 events per sorting gate (representative sorting gates are reported in **Fig. 2c** and **Supplementary** Fig. 10c). After seven days, cell pellets were collected, and DNA was extracted as detailed in the *High Molecular Weight DNA Extraction and Adenine Methylation* section.

### Scrambling OTX2-loxp6 cells

A slightly modified protocol was employed to scramble OTX2-loxp6 cells. One day prior to the experiment, 100,000 cells were seeded per well in a 24-well plate. To enhance Cre protein uptake and activity, the full medium was replaced with serum-free and PSG-free medium one hour before treatment. Purified TAT-Cre protein (Cambridge University Biochemistry Department) was added at concentrations of 1 µM and 500 nM, and the cells were incubated for 2 hours. After incubation, extracellular Cre protein was removed by treating the cells with TrypLE Express (ThermoFisher) for 10 minutes, followed by reseeding in full medium in 6-well plates. Two weeks later, the cells were harvested, stained with 10 µg/ml Hoechst 33342, and the fluorescence intensities of mScarlet in haploid G1 cells were analyzed using flow cytometry. Samples treated with 500 nM TAT-Cre were selected for further analysis due to their broader gene expression distributions, suggesting greater structural variant diversity. These cells were sorted into four gates based on mScarlet expression, with 10,000 events targeted per sorting gate (representative sorting gates are shown in **Supplementary** Fig. 2d). The sorted cells were cultured for an additional 14 days and then processed for high molecular weight DNA extraction, as described in the *High Molecular Weight DNA Extraction and Adenine Methylation* section.

### Flow cytometry

Flow cytometry analyses were conducted using a CytoFLEX Flow Cytometer (Beckman). Data acquisition was performed with CytExpert software, and subsequent analyses were carried out using FlowJo V10, CytoExploreR, and FCS Express 7. Events were initially gated to identify cells based on forward and side scatter properties. Singlets were then separated from doublets by evaluating the side scatter width and height. Finally, cells were analyzed for their respective fluorescence channels. Single-cell sorting for clones was performed using a Sony MA900, Sony SH800S, or MoFlo XDP cytometer.

### High molecular weight DNA extraction and adenine methylation

Between 2 and 4 million cells were collected by centrifugation, and high molecular weight (HMW) genomic DNA was extracted using the Monarch High Molecular Weight DNA Extraction Kit (New England BioLabs) with agitation speeds set to 1,500 rpm or 2,000 rpm. A modified version of the Monarch protocol was used to simultaneously assess DNA accessibility when required. In this modified protocol, 129 µl of nuclei prep buffer was combined with 10 µl of recombinant Cutsmart, 1 µl of 32 mM S-adenosylmethionine (final concentration 160 µM), 10 µl of 5000 Units/ml EcoGII methyltransferase (50 Units; New England BioLabs), and 5 µl of 20 mg/ml RNase A (200 µg; New England BioLabs). Approximately 2–3 million cells were resuspended in this reaction buffer and incubated at 37°C for 30 minutes. The reaction was then heated to 65°C for 5 minutes to inactivate the enzymes. Subsequently, 150 µl of nuclei prep buffer containing 10 µl of proteinase K was added to the nuclei preparation, and HMW DNA was extracted following the standard Monarch High Molecular Weight DNA Kit protocol. For samples containing six loxPsym sites obtained from screens, an additional step was included to eliminate low-molecular-weight DNA. This was achieved using the Short Fragment Eliminator Kit (Oxford Nanopore Technologies), following the manufacturer’s instructions.

### Cas9-targeted sequencing

Two crRNAs for each end of the enrichment region were designed using CHOPCHOP^61,62^ with the following settings: hg38 genome assembly, CRISPR/Cas9, and nanopore enrichment. Only protospacers meeting the criteria of efficiency scores > 50, GC content between 30% and 70%, self-complementarity = 0, and no off-target sites with one or two mismatches were selected. ALT-R tracrRNA and crRNAs were synthesised by Integrated DNA Technologies. The crRNAs used are listed in **Supplementary Table 1**. Cas9 enrichment was carried out using the Cas9 sequencing kit protocol with minor modifications (CAS9106 Protocol v109, Oxford Nanopore Technologies). Briefly, 5 µg of high molecular weight (HMW) DNA was dephosphorylated. tracrRNA and crRNA were annealed and complexed with HifiCas9 V3 (Integrated DNA Technologies) to form RNPs. The dephosphorylated DNA was incubated with Cas9 RNPs for 30 minutes at 37°C. Sequencing adapters were then ligated to the cleaved DNA, with the ligation step extended to 1 hour at room temperature. The libraries were purified using Ampure XP beads (Agilent), washed with Long Fragment Buffer, and eluted in an elution buffer for 16 hours. Sequencing was performed using the MinION Mk1B platform with R9.4.1 flow cells (FLO-MIN106).

### Long range PCR

To amplify the *OTX2* super-enhancer region for long-read sequencing, 250 ng of genomic DNA was used as input with 1 µM of forward and reverse primers. PCR amplification was performed using the LongAmp Hot Start Taq 2X Master Mix (New England BioLabs) under the following cycling conditions: initial denaturation at 94°C for 30 seconds, followed by 30 cycles of 94°C for 30 seconds, 59°C for 1 minute, and 65°C for 25 minutes, with a final extension at 65°C for 10 minutes. PCR products were purified using AMPure XP Beads (Beckman Coulter) at a bead-to-sample ratio of 0.66X and eluted with the elution buffer from the Monarch HMW DNA Extraction Kit for Cells & Blood. Genomic DNA from cell samples sorted into different gates was amplified using primers with custom index extensions for multiplexing (**Supplementary Table 1**). The resulting amplicons were sequenced on the MinION Mk1B platform using R10.4.1 flow cells (FLO-MIN114).

### RNA sequencing

RNA was extracted from 2–5 million flash-frozen cells using the RNeasy Plus Mini Kit (Qiagen). Libraries were prepared with the New England BioLabsNext® Ultra™ II Directional RNA Library Prep Kit for Illumina (New England Biolabs), multiplexed, and sequenced across two Illumina-HTP NovaSeq 6000 lanes, generating 150 bp paired-end reads. Transcript quantification was performed using Salmon and standard settings against a Salmon index constructed for the hg38 human reference genome. Further analysis was carried out with custom R scripts. Transcripts were aggregated to the gene level and TPMs calculated using `tximport`. Normalisation (rlog) and differential expression analysis were conducted using DESeq2^65^. Gene structure annotations were obtained from Ensembl release v110 and imported into R using biomaRt^66^. For all gene-related analyses, gene lists were filtered to include only those classified as "protein_coding," "lncRNA," "snRNA," "snoRNA," "miRNA," "rRNA," or "ribozyme," with expression base means (TPM) greater than 20.

### Base and variant calling

The Dorado basecaller with the dna_r9.4.1_e8_sup@v3.6 model was used for base calling without modifications, as well as for calling mCpG-modified bases. For m6A-modified base calling, the Guppy basecaller with the res_dna_r941_min_modbases-all-context_v001.cfg model was employed. The reads were aligned to the hg38 reference genome using either Dorado (for Cas9 sequencing) or bwa mem^67^ (for long-range PCR amplicon sequencing). BAM files were sorted and filtered using samtools, and further filtered for a region surrounding the OTX2 gene (chr14:56839500-56908200). Structural variants (used in **Fig. 2**, **Fig. 3**, and **Supplementary** Fig. 3-9) were called using SAVANA^68^ with relaxed QC settings (--buffer 100, --depth 1). Raw variants were filtered to include only those starting and ending within 50 bp of a loxPsym insertion site. The start and end positions of these variants were then adjusted to precisely align with the loxPsym insertion positions. Read names associated with the filtered variants were extracted. Variants were classified as either simple or complex: simple variants were defined as those with supporting reads plausibly having risen from a single rearrangement, and the rest were defined as complex. For experiments using the OTX2-loxp3 clonal cell line, reads supporting the wild type architecture were defined as those showing no rearrangements but displaying the expected number of loxPsym insertions. In experiments based on the OTX2-loxp6 design, WT reads were characterized as those with no rearrangements.

All identified architectures were validated by visually inspecting read coverage across the relevant domains. Read coverage was calculated using the `bamCoverage` function from deepTools with 100 nt windows. For architectures involving inversions, read coverage was calculated separately for the top and bottom strands, with negative values assigned to the bottom strand. The resulting coverage tracks were visually examined to confirm the accuracy of read grouping and genotype assignment.

### Calling structural variants

To optimise the detection and quantification of structural variants, we developed a custom caller, using the phasing of coverage and breakpoint signals at known loxPsym insertion sites to identify regulatory architectures. After base calling, a tiling of 300 bp short reads with a 200 bp overlap was extracted from each nanopore long read (oriented so that the unscrambled ends are on the forward strand), and aligned to the hg38 reference genome using bwa mem^67^ using parameters ‘-x ont2d’. This approach was used for **Fig. 4**, **Fig. 5**, and **Supplementary** Fig. 10-16.

For each long read, the following signals were extracted: (i) Coverage, calculated as the sum of mapped read bases across a domain, divided by the domain length. Based on this, the expected signal for a wild-type architecture is 3X, for a deletion event 0X, and for a duplication 6X. Coverage was recorded separately on both strands. (ii) Breakpoints, derived from tiled reads that overlap loxPsym sites and map discontinuously, with pairs of contiguous reads not overlapping. Deletion events are supported by contiguous read pairs that do not overlap and align to the same strand, inversion events are supported by contiguous read pairs that do not overlap but align to different strands, duplication events are supported by contiguous read pairs that do not overlap, align to the same strand, and have the second read aligning upstream of the first one, and inverted duplication events are supported by contiguous read pairs that do not overlap, align to different strands, and have the second read aligning upstream of the first one. (iii) LoxPsym sites, reported for any identified breakpoint when the loxPsym sequence (allowing for two mismatches) was found within the alignment. This approach iterates through all the tiled short reads for a given original long read to tally the different signals, and to assign a specific architecture. Reads where coverage and breakpoint signals were inconsistent, or where the loxPsym signal supporting the site at the position of rearrangement was absent, were discarded (about 18% of all reads).

Reads corresponding to the same architecture were grouped, and the architectures validated through visual inspection of read coverage across the domains of interest, as described above. High-confidence architectures generated from the scrambling of OTX2-loxp7 heterogeneous cell population, analyzed for their impact on *OTX2* expression, were defined as those supported by more than 15 reads and a difference of less than 25 from the averaged gene expression scores between duplicates (**Supplementary** Fig. 12a). Omitting the reproducibility filter, but retaining the coverage filter does not alter the conclusions of the modeling (see *Generalised linear modelling* section).

### Analysis of mCpG and m6A DNA modifications

Modified DNA bases were extracted from long-read data bam files with modified base calling information (see *Base and Variant Calling* above) using Fibertools^69^ as follows. The *`add-nucleosomes`* function was employed to annotate aligned reads with base modification data in the context of nucleosomes. The *`extract`* function was used to convert base modification information from BAM files into a tabular format. Individual reads were annotated with structural variant and sequencing gate information as determined previously (see *Base and Variant Calling* above). The reads were visualized in the R environment using ggplot2. DNase accessibility sites were identified from HAP1 DNase-seq data obtained from ENCODE (ENCFF289MVI), considering peaks with intensity > 2.5. The average adenine methylation for each read across DNase hypersensitivity sites was calculated and compared between structural variants using a two-tailed Student’s t-test.

### Quantifying impact of synthetic architectures on *OTX2* expression

After read grouping and genotype assignment, the distribution of reads supporting specific architectures across the dark, dim, medium, and bright sorting gates was analyzed to quantify the impact of each architecture on gene expression. Read counts and distributions for each experiment are detailed in **Supplementary File 1**. The relative frequency of each variant was determined by dividing the number of reads supporting that variant by the total number of reads within the sorting gate. Only architectures with sufficient coverage and/or reproducibility were included in the analysis as follows. For experiments using Cas9 sequencing, we considered variants observed in at least two sorting gates with a minimum of 25 total reads. For experiments using long-range PCR sequencing, we included variants observed in at least two sorting gates with a minimum of 15 total reads and a difference of less than 25 from the averaged gene expression scores between duplicates. To enable quantitative comparisons between architectures, gene expression scores were calculated as a weighted average of genotype frequencies across expression bins. We applied weights of -1, -0.5, 0.5, and 1 to the dark, dim, medium, and bright sorting gates, respectively. This results in scores of -1 for architectures with all reads in the dark gate, 1 for those with all reads in the bright gate, and 0 for architectures with reads evenly distributed across all four gates. The resulting scores were scaled to range from 0, representing the expression level of the largest deletion in a given experiment, to 100, representing wild-type expression, as references.

### Generalised linear modelling

Following an established method for fitting models of super-enhancer activity^11^, we trained generalised linear models that assume each constituent enhancer contributes additively, multiplicatively, or combinatorially to the expression level of *OTX2*. In this framework, the gene expression can be modelled using linear, exponential, or logistic functions based on super-enhancer activity, combined with a variable that accounts for various biological and experimental noise. The activity of the *OTX2* super-enhancer was modeled by an activity function 𝐴(𝑥) with 𝑥_𝑖_ as indicator variables for the presence of individual enhancer domain and β_𝑖_ as the coefficients that estimate the relative domain contributions to gene expression, such as 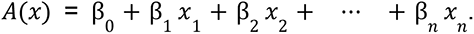 Each model predicts gene expression 𝑇(𝑥) as a monotonically increasing function of 𝐴(𝑥). The additive model assumes 𝑇(𝑥) = 𝐴(𝑥), the linear-exponential model 𝑇(𝑥) = 𝑒𝐴(𝑥) and the linear-logistic model 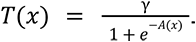 We considered alternative noise models that assume either normal or log-normal distributions. All models were fit in the *R* environment using the differential evolutionary algorithm package *DEoptim*^70^ and run for 10,000 steps. All replicates were considered simultaneously for model selection, and independently for quantification of enhancer domain contribution. The model with the best Bayesian Information Criterion was selected.

To evaluate the impact of coverage and the gene expression score difference thresholds used to define high-confidence architectures, we trained logistic-normal models as above using an increasing number of architectures ranked by coverage. For each group of architectures, five models were trained, and the best fits were selected to extract enhancer domain contributions (**Supplementary** Fig. 13g). When multiple models with similar good fits were generated, enhancer contributions were averaged. Omitting the reproducibility filter retained the large effect of R5-2 at high coverages, and that including architectures observed in only one replicate, or not consistently quantified in two replicates at high coverage (for which measurement noise cannot easily explain the discrepancy) induced noise in model fits, motivating the use of both coverage and reproducibility filters.

### Enhancer/promoter interaction and Gene expression predictions of deletions

Connections between the *OTX2* promoter and enhancer candidates were first predicted with the Activity-By-Contact (ABC) model^42^ using available genomic data for DNA accessibility (ENCODE accessions: ENCFF303JZI, ENCFF427IVB), H3K27ac ChIP-seq (ENCODE accessions: ENCFF496HYU, ENCFF806PAF) and Hi-C contacts (ENCODE accession: ENCFF898HRO, using a resolution of 5000 bp). The ABC model was run using default parameters other than the input datasets, and the recommended threshold of an ABC score of 0.027 was used to filter predicted enhancer-promoter contacts. In a second instance, we used pre-computed predictions (ENCODE accession: ENCFF331ONH) by the refined ENCODE-rE2G model^42,43^, which was previously trained on CRISPRi data from the K562 cell line to predict interactions in HAP1 cells using HAP1 DNA accessibility data. To predict gene expression using Borzoi, one-hot encoded 524,288 bp sequences containing wildtype and deletion variants of the *OTX2* gene and enhancer cluster were fed to the four Borzoi model folds, of which we took the average across models. We extracted the average predictions per 32 bp bin and for each tissue took the mean of the predictions made using the prediction heads associated with that tissue. Deletion variant sequences were generated by deleting the appropriate parts of the sequence and concatenating further genomic sequence on the right-hand side to create full-length (524 kb) sequences. Expression values for the *OTX2* gene were calculated by summing the expression over the window chr14:56,799,905-56,810,479, which fully covers the *OTX2* gene and no other genes. Borzoi model code and weights were downloaded by following the guide at https://github.com/calico/borzoi.

### Transcription factor binding sites

Transcription factors (TFs) associated with the R5-2 enhancer domain were identified from ATAC-seq signals using maxATAC^45^ as follows. Publicly available ATAC-seq data^71^ for HAP1 cells (accession numbers SRR6410048 and SRR6410049) were downloaded from the Sequence Read Archive (SRA) via the SRA toolkit. Reads from both duplicates were combined and aligned to the human reference genome (hg38) using bwa mem^67^. The ATAC-seq signal was normalized using the `prepare` function in maxATAC. Peaks were identified from the normalized signal using the `bdgpeakcall` function in MACS2^72^ with the following parameters: cutoff of 0.1, max-gap of 100, and minimum length of 50. Peaks were further filtered to include only those with widths between 500 bp and 10 kb and scores exceeding 250.

To assess TF binding at the *OTX2* super-enhancer, the `predict` function of maxATAC was used, analysing the binding of 135 TFs as follows. TFs of interest were selected based on the probability (maxATAC score) and selectivity of their binding to the single R5-2 ATAC-seq peak. Binding selectivity was calculated as the fraction of the average maxATAC score at the R5-2 peak from the sum across all 10 peaks within the *OTX2* super-enhancer. Five TFs, *FOXA1*, *FOXK2*, *YY1*, *RFX5*, and *MYB*, were identified as expressed in HAP1 cells (≥ 10 RPKM) and demonstrating high selectivity and/or probability of binding (0.4 selectivity and/or binding score threshold) within the R5-2 domain. Consensus DNA binding sequences for these TFs were identified using the `searchSeq` function from the R/Bioconductor TFBSTools package^73^, with position weight matrices sourced from Jaspar (https://jaspar.elixir.no/) and a minimum score threshold of 60%. This analysis revealed binding sites for FOXA1 (6 sites), YY1 (1 site), RFX5 (1 site), and MYB (4 sites). Binding sites for FOXK2 overlapped with those identified for FOXA1.

### Deleting transcription factor binding sites

Short deletions of 6 to 8 bp were targeted at selected transcription factor binding sites using prime editing. epegRNAs were designed with PRIDICT2.0^74^ and cloned as described above. The first nucleotide of the spacer was adjusted to be a G for better U6 expression. The epegRNA sequences are detailed in **Supplementary Table 2**. Two rounds of transfection were conducted in triplicate as described above. Between 1 and 2 million cells were harvested for FACS sorting, selecting the ∼5% of cells exhibiting lower OTX2-mScarlet expression. The cells were allowed to recover and expand for a week before undergoing a second round of FACS sorting. Subsequently, the cells were expanded to prepare cell pellets for gDNA extraction as described above. The R5-2 region of interest was amplified using primers listed in **Supplementary Table 1** and prepared for Illumina sequencing with index adapters. Amplicons were sequenced on a MiSeq system using 300 paired-end cycles. After demultiplexing, paired-end reads were merged using PEAR^75^. Reads containing both forward and reverse PCR primer sequences were identified using the amplicon function of SeqKit^76^, allowing for up to four mismatches. Amplicon sequences were then analyzed to identify and quantify deletions using CRISPResso2^77^. CRISPResso2 was used in ‘Prime Editing’ mode, with spacer and extension sequences provided to identify intended edits. In ‘Standard’ mode, the `--quantification_window_coordinates` parameter was used to specify the positions of targeted sites and to quantify any deletions overlapping transcription factor binding sites (TFBSs) of interest. Deletion frequencies were calculated as the ratio of reads reporting a deletion overlapping a given TFBS to the total number of reads analyzed for a given condition. Deletion enrichments were computed by dividing the values obtained for each replicate by the averaged values observed for amplicons from unsorted cells.

## Data availability

Sequencing data will be made available at Gene Expression Omnibus (GEO). All other data are available in the article and its Supplementary Information (Supplementary Figures 1-16, Supplementary Tables 1-2), Dataset 1 (read counts supporting individual architectures in each experiment and replicate), or from the corresponding author upon request.

## Code availability

The Bash, R and Python code used to analyse the data and generate the figures reported in the manuscript are available at Github (https://github.com/pierre-murat/escramble_OTX2).

## Software

Genomics: bedtools (2.29.0), benchling (accessed between 2019-2024), bwa (0.7.17), ChromHMM (1.24), CRISPResso2 (46), deeptools (3.5.3), Dorado (0.7.2), fibertools (0.3.2), guppy (6.3.8 and 6.4.6), IGV (2.16.2), macs2 (2.2.9.1), maxATAC (1.0.6), minimap2 (2.22), modbam2bed (0.9.5), modkit (0.1.12), mosdepth (0.3.3), PEAR (0.9.6), R (4.1.3), salmon, Salmon (2.0.0.55), samtools (1.14), SAVANA (1.0.0), seqkit (0.10.1), nanomonsv (0.6.0). Flow cytometry: FlowJo (v10), CytoExploreR (1.1.0). CytExpert.

R packages: BSgenome.Hsapiens.UCSC.hg38 (1.4.5), biomaRt (2.50.3), Biostrings (2.74.1), DESeq2 (1.34.0), DEoptim (2.2-8), dplyr (1.1.4), fgsea (1.20.0), forcats (1.0.0), ggplot2 (3.5.1), ggrepel (0.9.3), ggpointdensity (0.1.0), plotgardnerer (1.4.2), RBioinf (1.48.0), reversetranslate (1.0.0), rtracklayer (1.66.0), scales (1.3.0), shortRead (1.46.0), spgs (1.0-3), stringr (1.5.1), StructalVariantAnnotation (1.10.1), tidyverse (1.3.2), TFBSTools (1.44.0), tximport (1.22.0), viridis (0.6.2), wesanderson (0.3.6).

Python packages: Levenshtein (0.26.1), matplotlib (3.9.2), numpy (2.1.3), pandas (2.2.3), pysam (0.22.1), seaborn (0.13.2).

## Notes

### Competing Interest Statement

L.P. receives remuneration and stock options from ExpressionEdits.

