## Supplementary Information for "Resolution of a human super-enhancer by targeted genome randomisation"

### Supplementary Figures

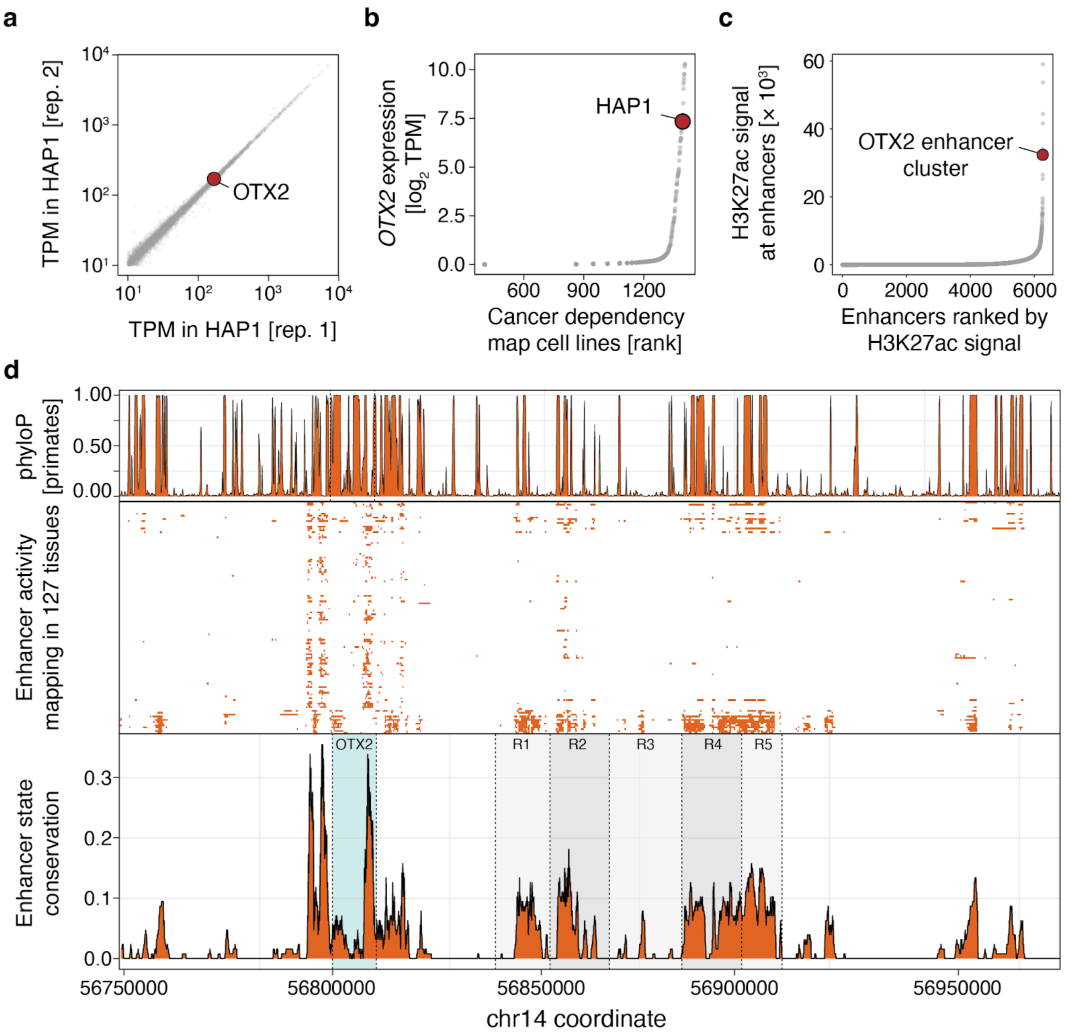


**Supplementary Figure 1. Structure and expression of the *OTX2* locus in HAP1 cells. (a)** Expression level (TPM; x- and y-axis) of genes (markers) in two replicate measurements of HAP1 cells. Red marker: *OTX2*. **(b)** *OTX2* expression ($log_2$TPM, y-axis) in 1,406 cancer cell lines (points) sourced from the Cancer Cell Line Encyclopedia (CCLE) Data Portal (22Q2 release, https://sites.broadinstitute.org/ccle/datasets) ranked by *OTX2* expression (x-axis). The HAP1 cell line is marked in red. **(c)** H3K27ac ChIP-seq signal (y-axis) at 'stitched' enhancers ranked by input-normalised H3K27ac ChIP-seq signal (x-axis). **(d)** Enhancer sequence conservation (top panel), activity (middle panel), and activity conservation (bottom panel) values (y-axis) across the *OTX2* locus in HAP1 cells (x-axis), mapped using the core 15-state ChromHMM model across 127 cell types from the Roadmap Epigenomics dataset (https://egg2.wustl.edu/roadmap/web_portal/chr_state_learning.html). *OTX2* and the R1-R5 domains are highlighted in green and grey boxes, respectively.

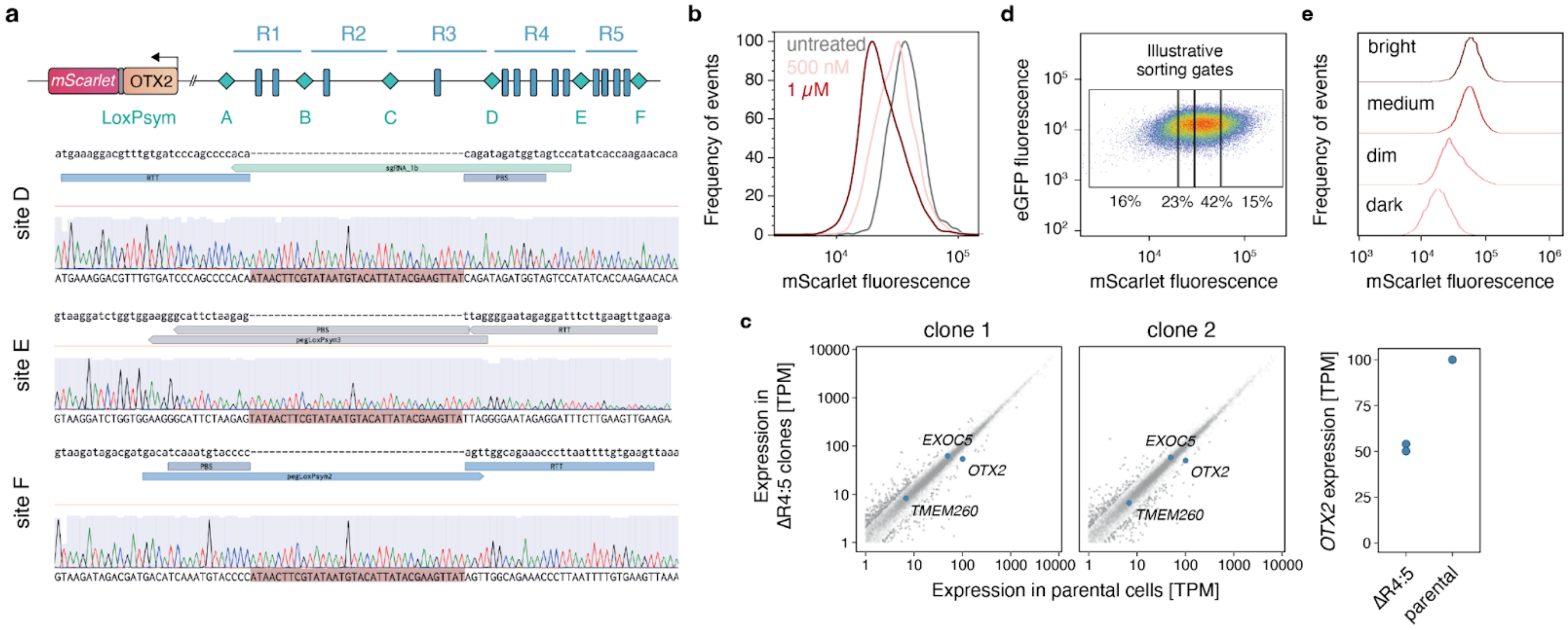


**Supplementary Figure 2. Engineering and scrambling of the OTX2-loxp3 and loxp6 cell lines. (a)** Genotyping of OTX2-loxp3 clonal cells showing precise insertion of loxPsym at targeted sites. **(b)** Frequency (y-axis) of OTX2-loxp6 cells with mScarlet fluorescence (x-axis) before (grey) and after scrambling (pink:500 nM Cre, red: 1 μM Cre). **(c)** Loss of the R45 super-enhancer domain affects *OTX2* expression. Left: Gene expression (x- and y- axis) in two ΔR45 clones (panels) compared to the parental cell line (y-axis). Highlights: *OTX2* and the two adjacent genes, *EXOC5* and *TMRM260*. Right: *OTX2* expression (y-axis) in ΔR45 clones and the parental cell line (x-axis). **(d)** Diagram of sorting gates (boxes) used to isolate OTX2-loxp6 cells with varying levels of *OTX2* expression. **(e)** Frequency of cells (y-axis) with different mScarlet fluorescence (x-axis) of scrambled OTX2-loxp3 clonal cells 20 days post-scrambling and sorting into bins (panels)..

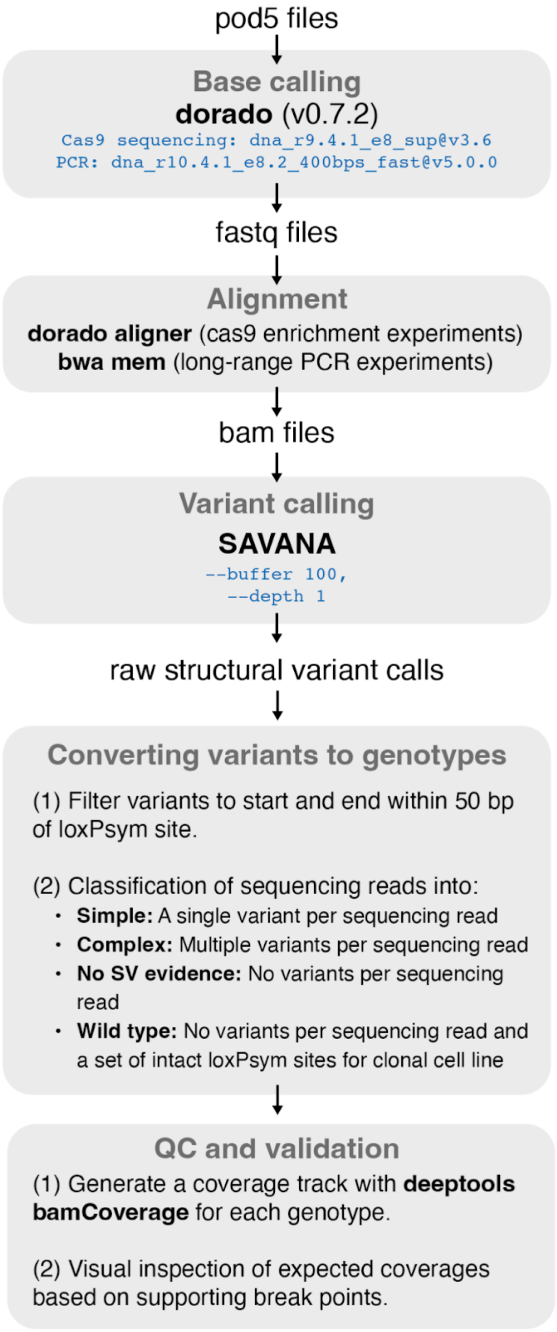


**Supplementary Figure 3. Pipeline to genotype sequencing reads.** Schematic outlining the bioinformatics pipeline used to process sequencing data and call Cre-induced structural variants (used for Figures 2 and 3, Supplementary Figures 1-9; Methods). pod5 files were base called using dorado and aligned with dorado aligner. SAVANA was used with relaxed parameters to call raw structural variants which were filtered based on proximity to loxPsym insertion sites. Sequencing reads are then classified into simple, complex, no SV evidence, or wild type based on read names and variant information. Finally, called genotypes were validated by visual inspection and correction, if needed, from read coverage tracks.

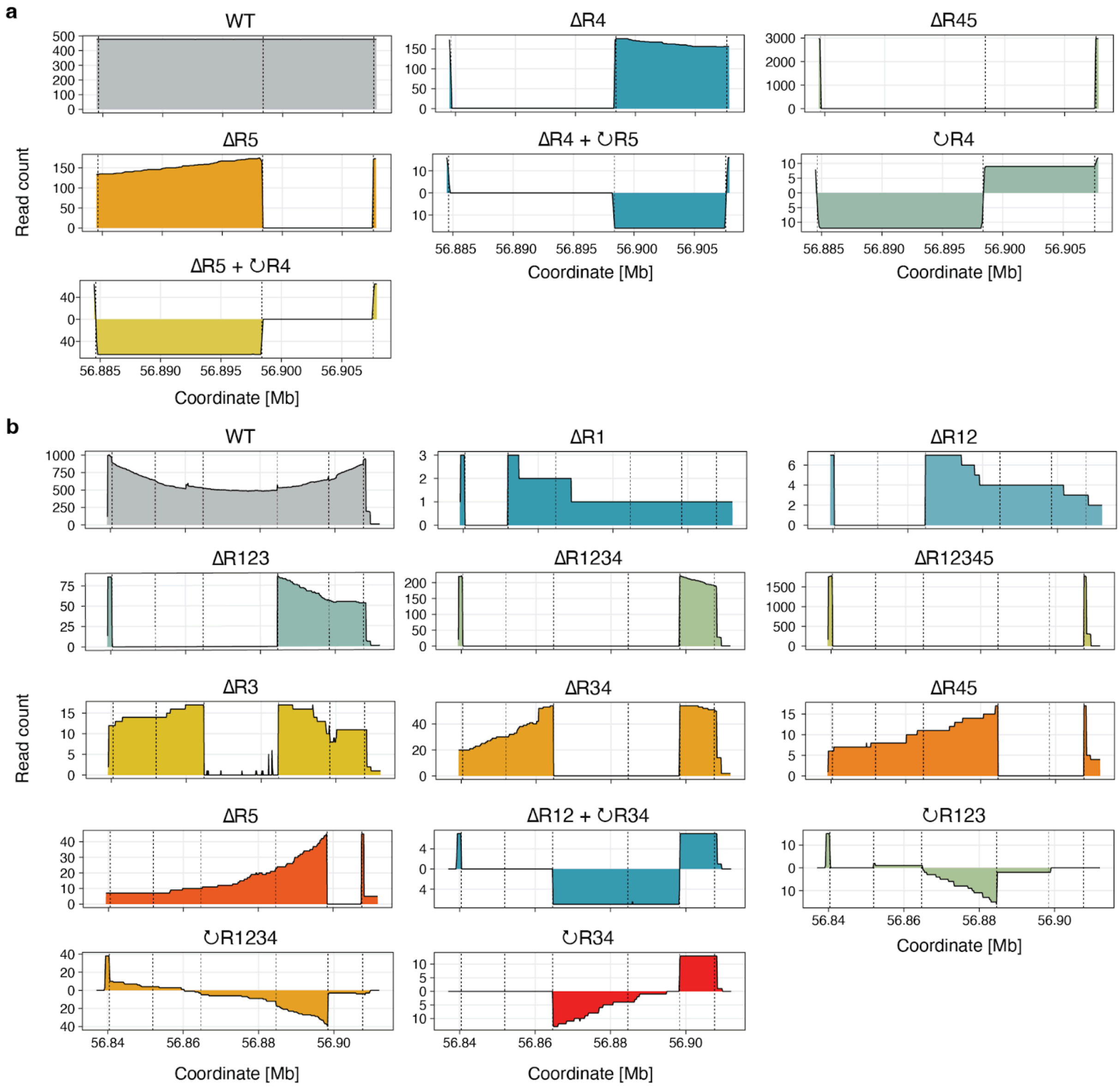


**Supplementary Figure 4. Synthetic architectures identified through scrambling of clonal cell lines.** After recombination at the engineered *OTX2* locus, followed by cell sorting, Cas9 enrichment, and sequencing, reads supporting specific genotypes were grouped (**Methods**), leading to the identification of 21 unique architectures. (**a**) Read coverage (y-axis) across the region (x-axis), on the forward strand (positive values) and bottom strand (negative values) for architectures identified from scrambled OTX2-loxp3 cells. Dashed lines: locations of the three clonal loxPsym sites. (**b**) As (a), but for OTX2-loxp6 cells.

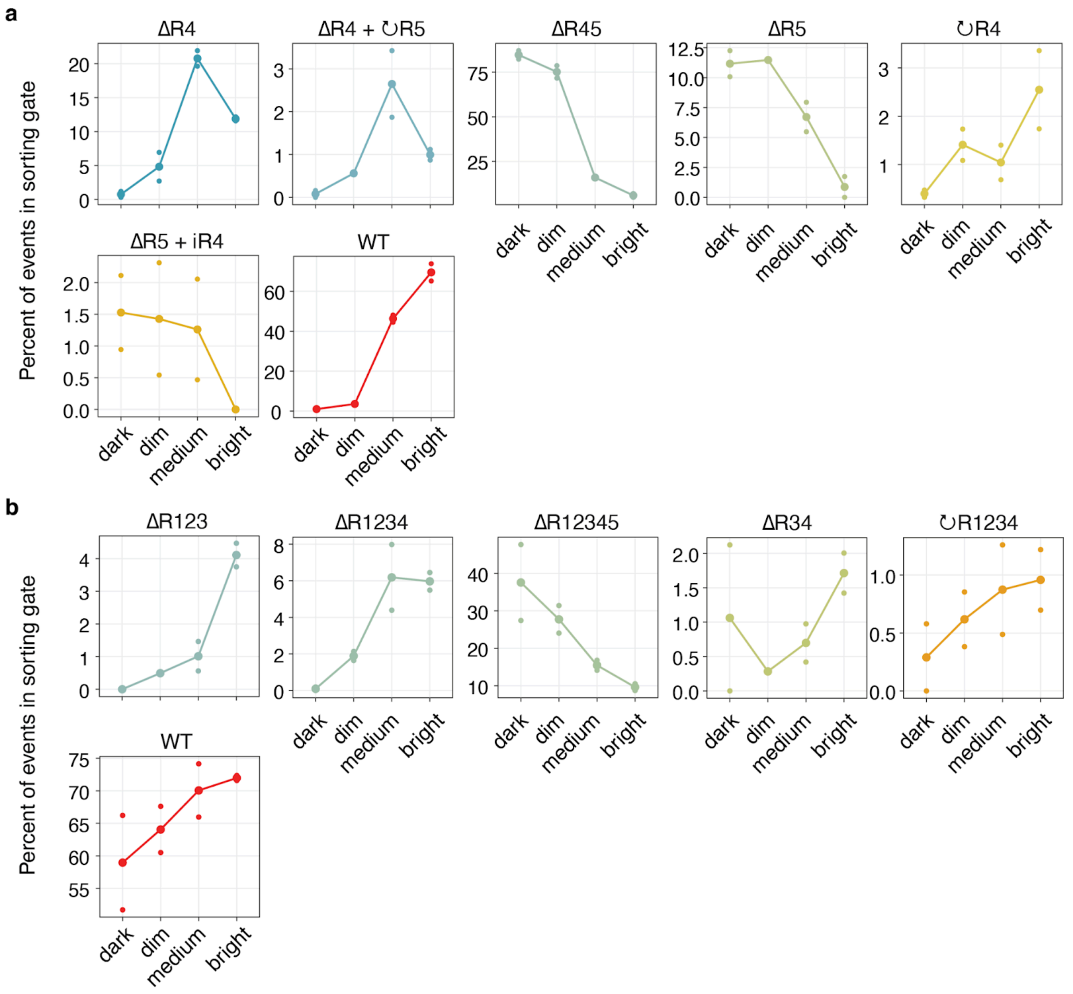


**Supplementary Figure 5. Impact of synthetic architectures identified from the scrambling of clonal cell lines on OTX2 expression.** Distribution of reads associated with architectures with consistent read coverage from the (**a**) Percentage of reads (y-axis) associated with the identified architectures (panels) from the scrambled OTX2-loxp3 cells within the sorting gates (x-axis). Markers: replicate values. (**b**) As (a), but for OTX2-loxp6 cells. Sorting gates are defined in **Fig. 2c** and **Supplementary Fig. 2d**.

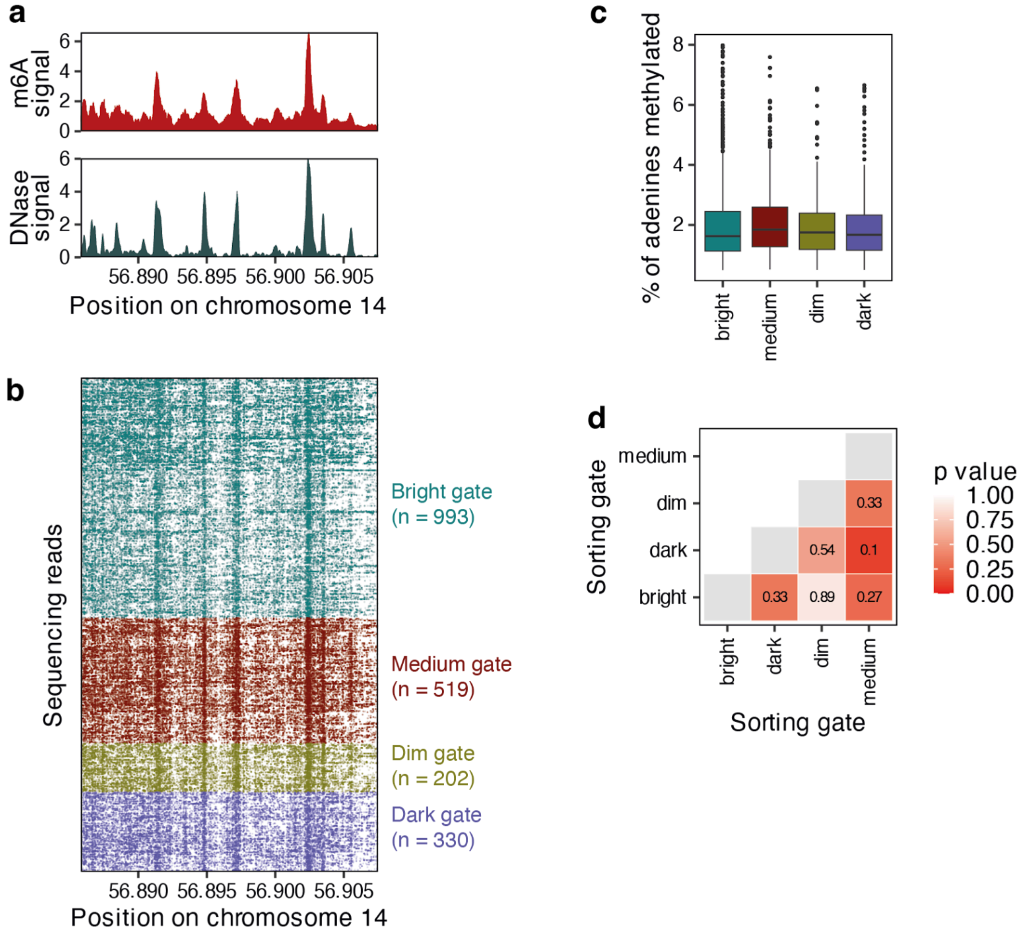


**Supplementary Figure 6. DNA accessibility is consistent across sorting gates and technologies. (a)** Comparison of inferred accessibility (y-axis) from Fiber-seq (dark red, top panel) and a publicly available DNase-seq data set (ENCFF314TDG, dark green, bottom panel) for a region of chromosome 14 covering the *OTX2* super-enhancer (x-axis). The m6A data represents all wild-type reads across OTX2-loxp3 and OTX2-loxp6 cells smoothed with a rolling window of size 100. **(b)** Individual nanopore sequencing reads (rows) with m6A-modified bases (dashes) for wildtype reads across sorting gates (colors). n: number of sequencing reads for each sorting gate. **(c)** Percentage of methylated adenines (y-axis) for different sorting gates (colors, x-axis). Box plots: medians and interquartile ranges. **(d)** Pairwise tests of mean difference between all sorting gates (x- and y-axes) colored and labelled by Benjamini-Hochberg adjusted *P* values from two-tailed Student’s t-tests.

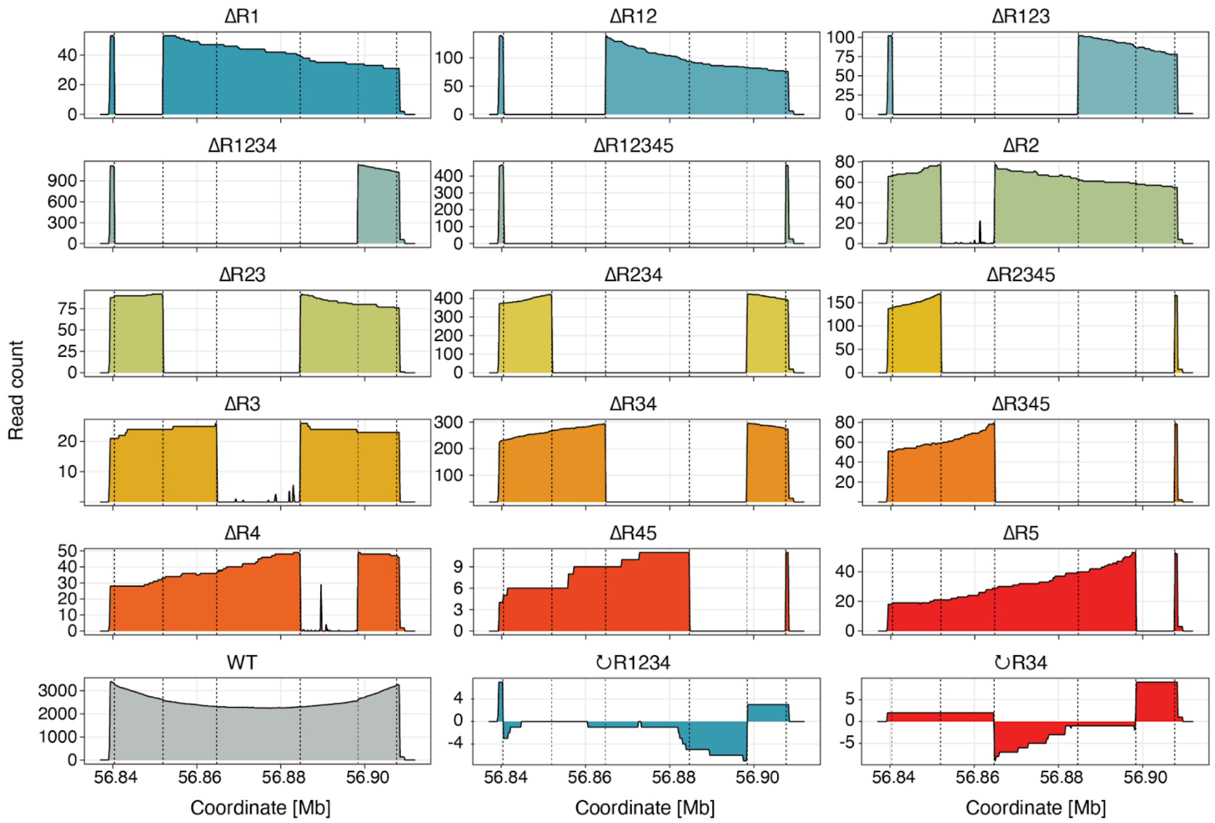


**Supplementary Figure 7. Synthetic architectures identified through scrambling of an heterogeneous OTX2-loxp6 cell population.** To increase the diversity and representation of structural variants, a pooled epegRNA transfection approach was used to insert six loxPsym sites, following the design outlined in **Fig. 2a**. After recombination at the engineered OTX2 locus in the diverse cell population, cell sorting, Cas9 enrichment coupled with long-read sequencing was used to identify 18 unique architectures (**Methods**). Read coverage (y-axis) across the region (x-axis), on the forward strand (positive values) and bottom strand (negative values). Dashed lines: locations of the three clonal loxPsym sites.

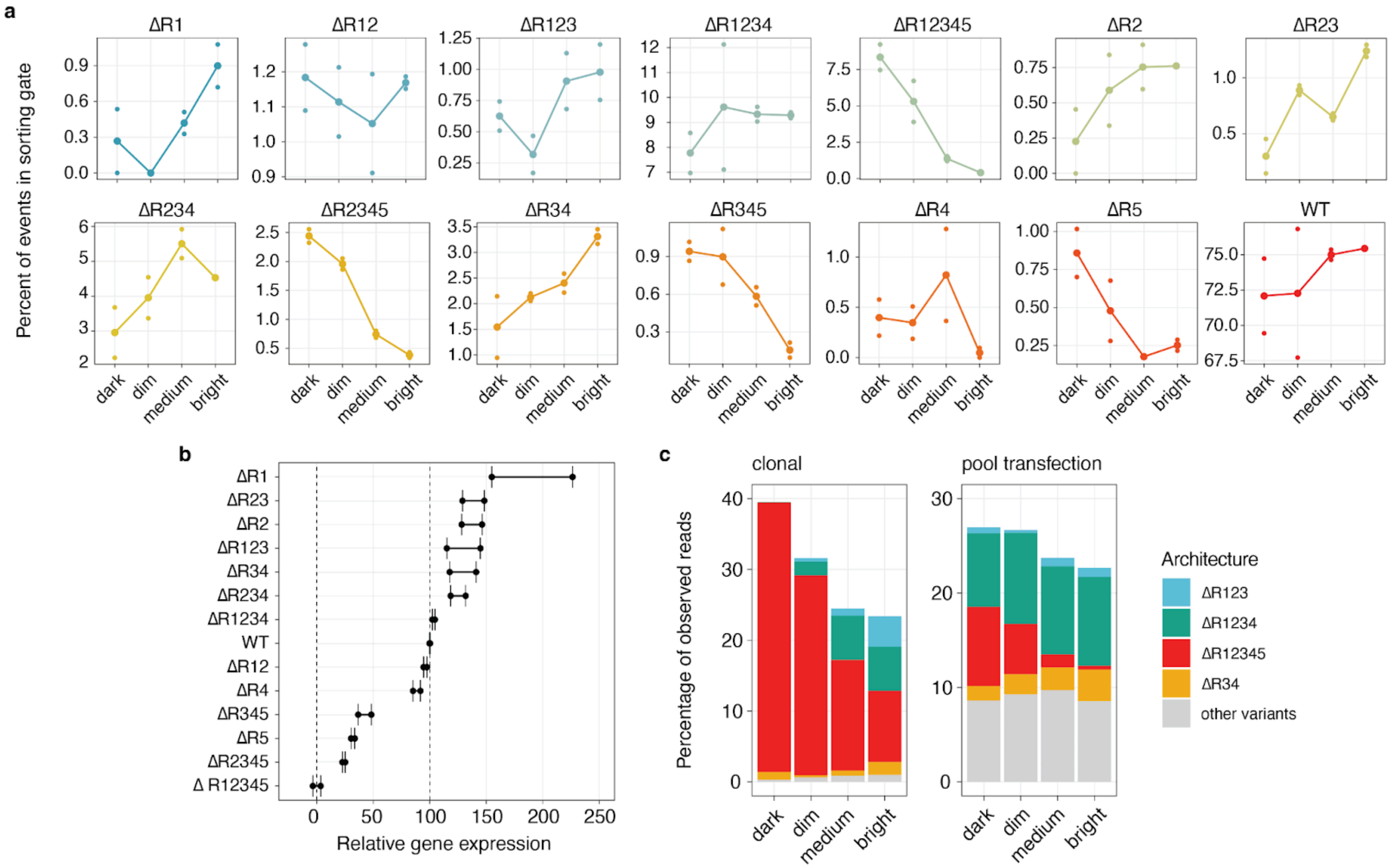
a
Percent of events in sorting gate
ΔR1
ΔR12
ΔR123
ΔR1234
ΔR12345
ΔR2
ΔR23
ΔR234
ΔR2345
ΔR34
ΔR345
ΔR4
ΔR5
WT
dark
dim
medium
bright
b
ΔR1
ΔR23
ΔR2
ΔR123
ΔR34
ΔR234
ΔR1234
WT
ΔR12
ΔR4
ΔR345
ΔR5
ΔR2345
Δ R12345
0
50
100
150
200
250
Relative gene expression
c
clonal
pool transfection
Percentage of observed reads
dark
dim
medium
bright
Architecture
ΔR123
ΔR1234
ΔR12345
ΔR34
other variants

**Supplementary Figure 8. Impact on OTX2 expression and diversity of synthetic architectures resulting from scrambling a heterogeneous OTX2-loxp6 cell population.** (**a**) Percentage of reads (y-axis) associated with the identified architectures (panels) within the sorting gates (x-axis). Markers: replicate values. (**b**) Relative gene expression (x-axis) for different architectures (y-axis). Markers: replicate values. Dashed lines: 0 (largest deletion) and 100 (wild type). (**c**) Percentage of observed reads (bar height) for different architectures (colors) derived from scrambling clonal (left) or heterogeneous (right) OTX2-loxp6 cell populations. Contributions from WT architectures are excluded for clarity.

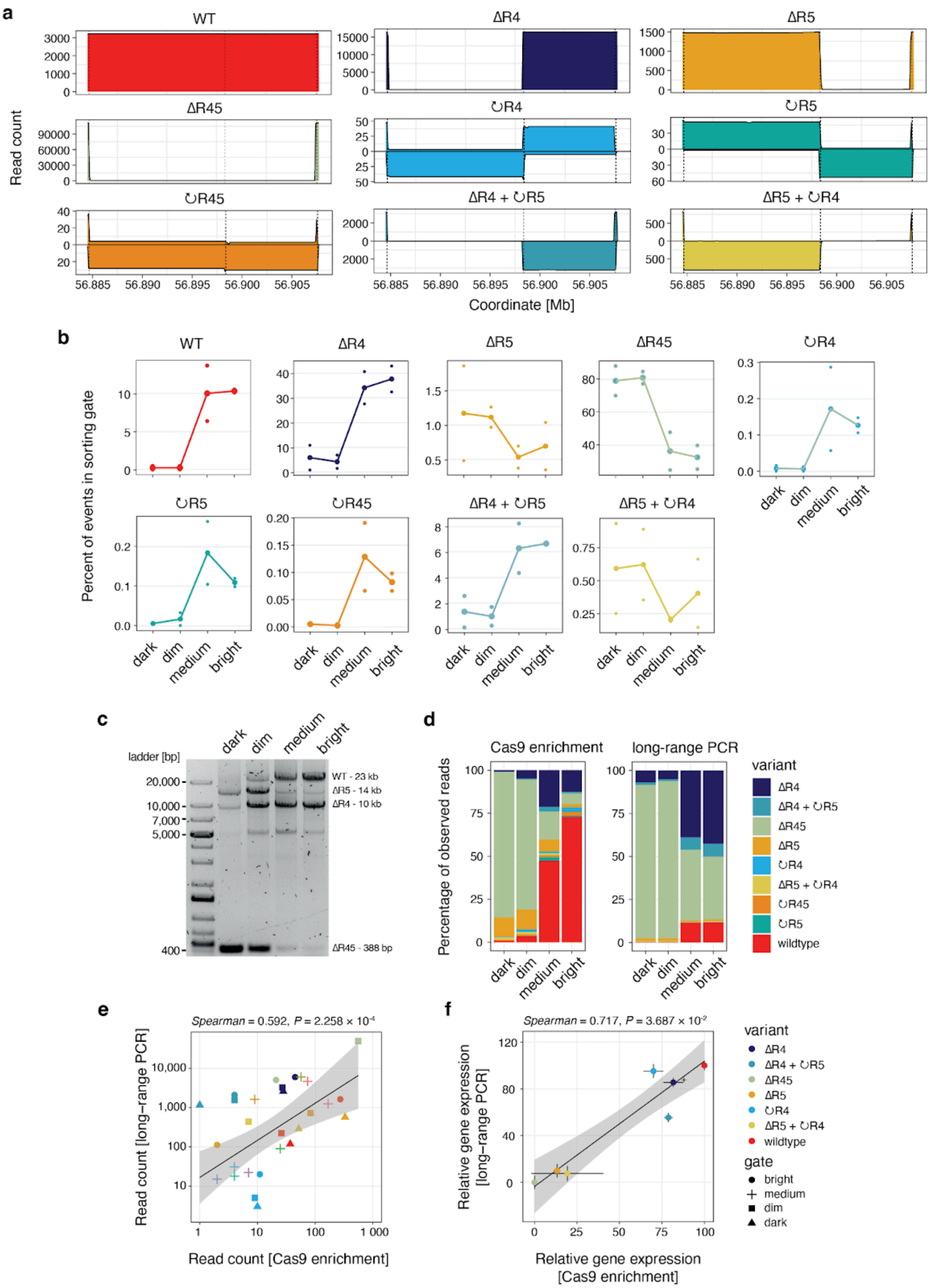


**Supplementary Figure 9. Comparison of Cas9 enrichment and long-range PCR approaches for characterizing scrambled OTX2-loxp3 cells.** Following recombination at the engineered *OTX2* locus in OTX2-loxp3 cells and subsequent cell sorting, long-range PCR was used to amplify the R45 super-enhancer domain (**Methods**). Sequencing and structural variant analysis identified all 9 possible architectures, with sufficient coverage to evaluate their effects on OTX2 expression. **(a)** Read coverage (y-axis) across the R45 super-enhancer region (x-axis), on the forward strand (positive values) and bottom strand (negative values). Dashed lines: locations of the three clonal loxPsym sites. **(b)** Percentage of reads (y-axis) associated with the identified architectures (panels) within the sorting gates (x-axis). Markers: replicate values. **(c)** Structural variant diversity visualised through gel electrophoresis of the sequencing library, illustrating the distribution of structural variants across sorting gates. (**d**) Percentage of reads (bar height) from each architecture (colors) associated with each sorting gate (x-axis) observed using the Cas9 enrichment approach (left) and the long-range PCR method (right). **(e)** Coverage of individual variants (colors) in different sorting gates (markers) using long-range PCR sequencing (y-axis) and Cas9-enrichment (x-axis). **(f)** Relative gene expression of individual variants (colors) using long-range PCR sequencing (y-axis) and Cas9-enrichment (x-axis). Points and whiskers represent means and ranges respectively.

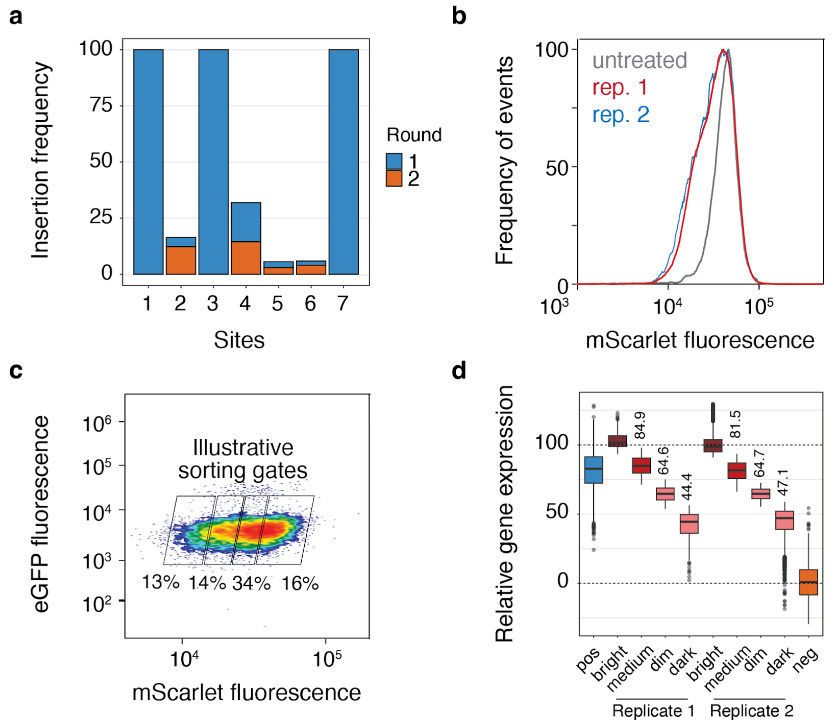


**Supplementary Figure 10. Characterization of the OTX2-loxp7 heterogeneous cell population. (a)** The OTX2-loxp7 heterogeneous cell population was derived from the OTX2-loxp3 cell line, which contains clonal insertions of loxPsym sequences at sites 1, 3, and 7. Insertion frequency (y-axis) after one (blue) and two (orange) rounds of co-transfection using a pool of four epegRNAs at all seven sites (x-axis). **(b)** Frequency (y-axis) of mScarlet fluorescence intensity (x-axis) in the OTX2-loxp7 cell populations. Grey: untreated; red: replicate 1; blue: replicate 2. **(c)** Exemplar sorting gates (outlined boxes) used for isolating cells (markers) based on varying levels of OTX2-mScarlet fluorescence (x-axis), with eGFP fluorescence (y-axis) serving as a control. Color: density of cells. **(d)** Relative gene expression (y-axis) for cells from the four sorting gates (x-axis), and each replicate (left and right half) after normalization. Unperturbed cells (pos) are included for comparison. Normalization was performed by subtracting the mean of the mScarlet-negative cells (neg) and dividing each individual value by the mean of the bright gates. Mean values are shown above the boxes.

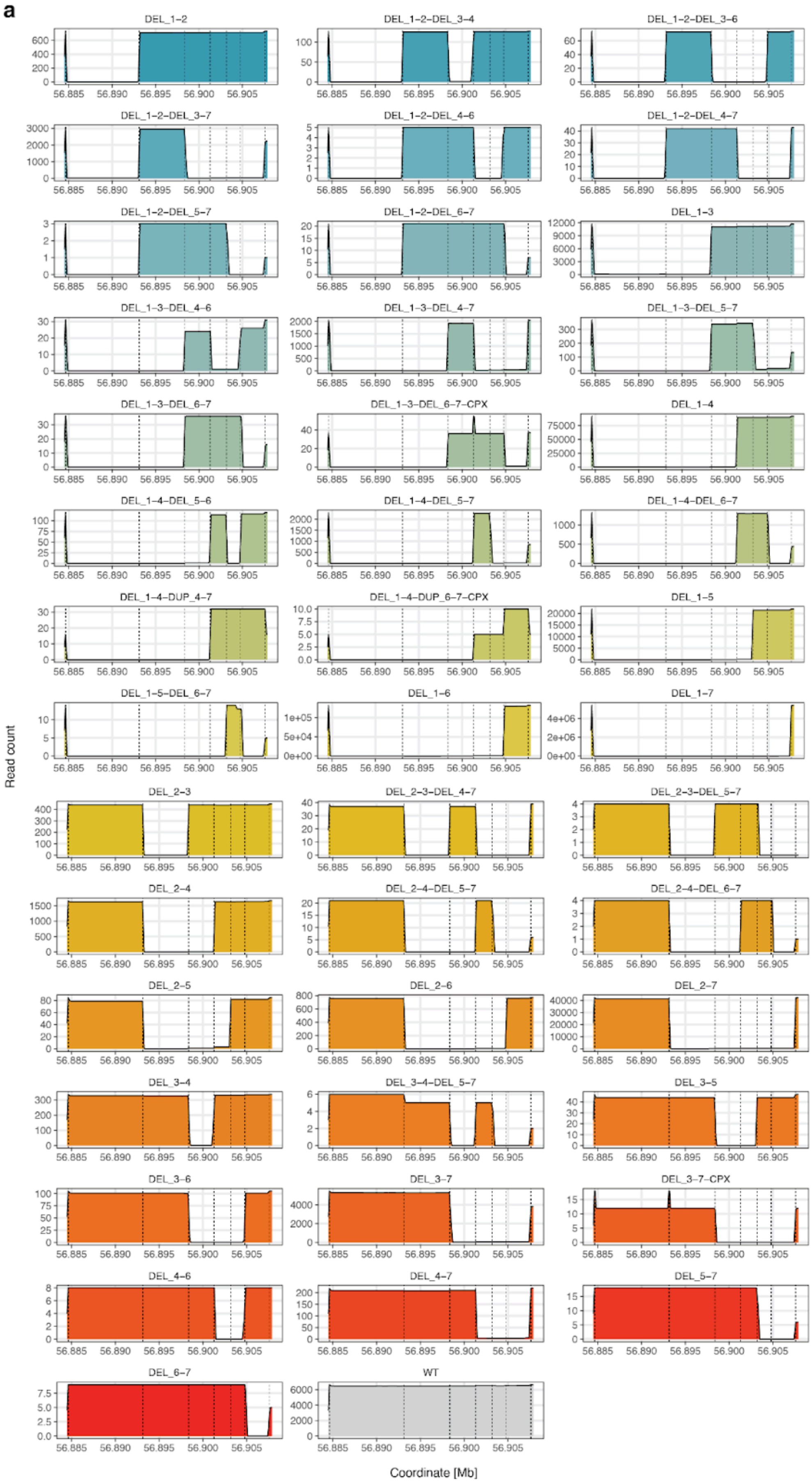
a
DEL_1–2
DEL_1–2–DEL_3–4
DEL_1–2–DEL_3–6
DEL_1–2–DEL_3–7
DEL_1–2–DEL_4–6
DEL_1–2–DEL_4–7
DEL_1–2–DEL_5–7
DEL_1–2–DEL_6–7
DEL_1–3
DEL_1–3–DEL_4–6
DEL_1–3–DEL_4–7
DEL_1–3–DEL_5–7
DEL_1–3–DEL_6–7
DEL_1–3–DEL_6–7–CPX
DEL_1–4
DEL_1–4–DEL_5–6
DEL_1–4–DEL_5–7
DEL_1–4–DEL_6–7
DEL_1–4–DUP_4–7
DEL_1–4–DUP_6–7–CPX
DEL_1–5
DEL_1–5–DEL_6–7
DEL_1–6
DEL_1–7
DEL_2–3
DEL_2–3–DEL_4–7
DEL_2–3–DEL_5–7
DEL_2–4
DEL_2–4–DEL_5–7
DEL_2–4–DEL_6–7
DEL_2–5
DEL_2–6
DEL_2–7
DEL_3–4
DEL_3–4–DEL_5–7
DEL_3–5
DEL_3–6
DEL_3–7
DEL_3–7–CPX
DEL_4–6
DEL_4–7
DEL_5–7
DEL_6–7
WT
Read count
Coordinate [Mb]

**b**

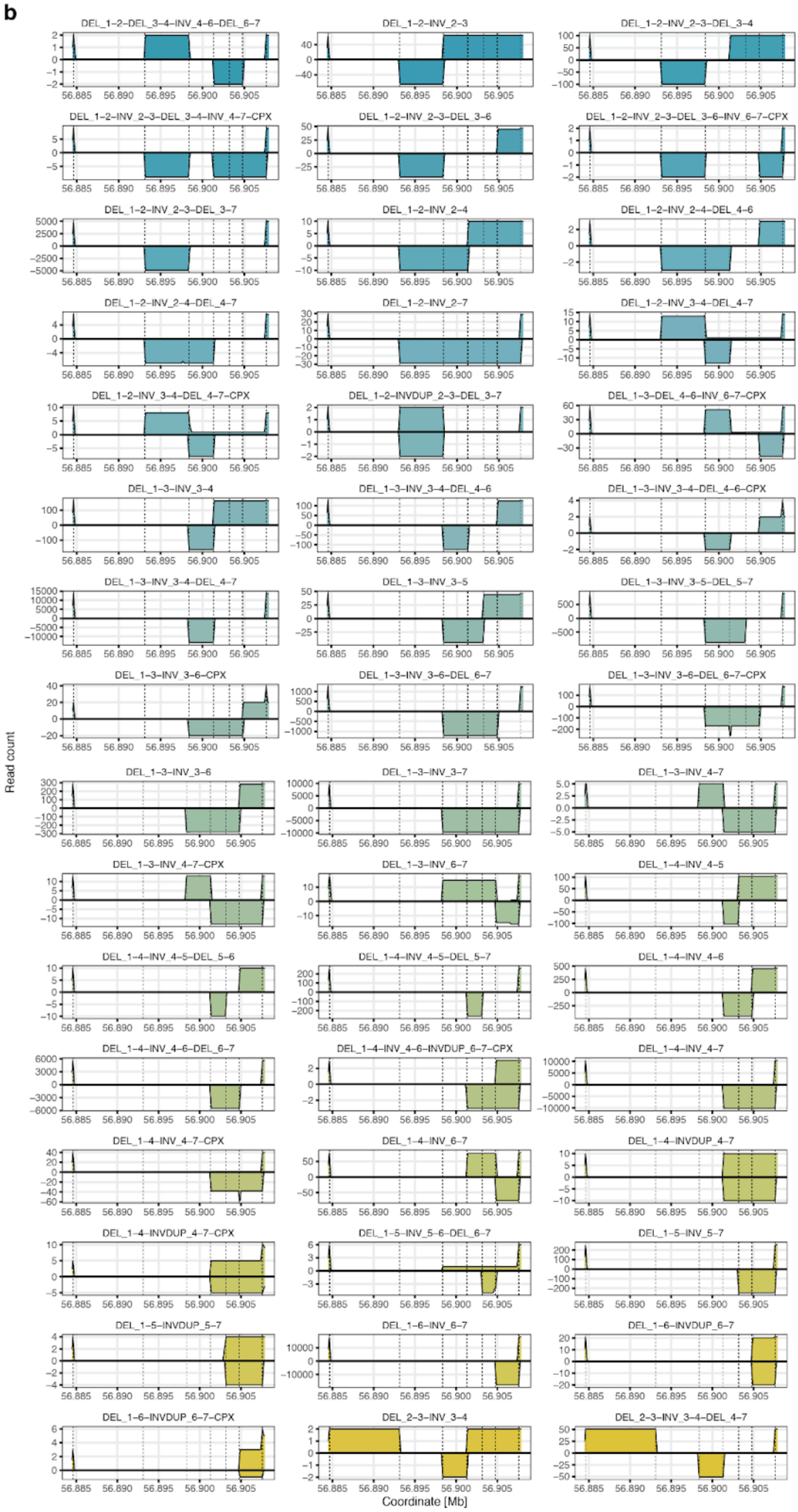
DEL_1-2-DEL_3-4-INV_4-6-DEL_6-7
DEL_1-2-INV_2-3
DEL_1-2-INV_2-3-DEL_3-4
DEL_1-2-INV_2-3-DEL_3-4-INV_4-7-CPX
DEL_1-2-INV_2-3-DEL_3-6
DEL_1-2-INV_2-3-DEL_3-6-INV_6-7-CPX
DEL_1-2-INV_2-3-DEL_3-7
DEL_1-2-INV_2-4
DEL_1-2-INV_2-4-DEL_4-6
DEL_1-2-INV_2-4-DEL_4-7
DEL_1-2-INV_2-7
DEL_1-2-INV_3-4-DEL_4-7
DEL_1-2-INV_3-4-DEL_4-7-CPX
DEL_1-2-INVDUP_2-3-DEL_3-7
DEL_1-3-DEL_4-6-INV_6-7-CPX
DEL_1-3-INV_3-4
DEL_1-3-INV_3-4-DEL_4-6
DEL_1-3-INV_3-4-DEL_4-6-CPX
DEL_1-3-INV_3-4-DEL_4-7
DEL_1-3-INV_3-5
DEL_1-3-INV_3-5-DEL_5-7
DEL_1-3-INV_3-6-CPX
DEL_1-3-INV_3-6-DEL_6-7
DEL_1-3-INV_3-6-DEL_6-7-CPX
DEL_1-3-INV_3-6
DEL_1-3-INV_3-7
DEL_1-3-INV_4-7
DEL_1-3-INV_4-7-CPX
DEL_1-3-INV_6-7
DEL_1-4-INV_4-5
DEL_1-4-INV_4-5-DEL_5-6
DEL_1-4-INV_4-5-DEL_5-7
DEL_1-4-INV_4-6
DEL_1-4-INV_4-6-DEL_6-7
DEL_1-4-INV_4-6-INVDUP_6-7-CPX
DEL_1-4-INV_4-7
DEL_1-4-INV_4-7-CPX
DEL_1-4-INV_6-7
DEL_1-4-INVDUP_4-7
DEL_1-4-INVDUP_4-7-CPX
DEL_1-5-INV_5-6-DEL_6-7
DEL_1-5-INV_5-7
DEL_1-5-INVDUP_5-7
DEL_1-6-INV_6-7
DEL_1-6-INVDUP_6-7
DEL_1-6-INVDUP_6-7-CPX
DEL_2-3-INV_3-4
DEL_2-3-INV_3-4-DEL_4-7
Read count
Coordinate [Mb]

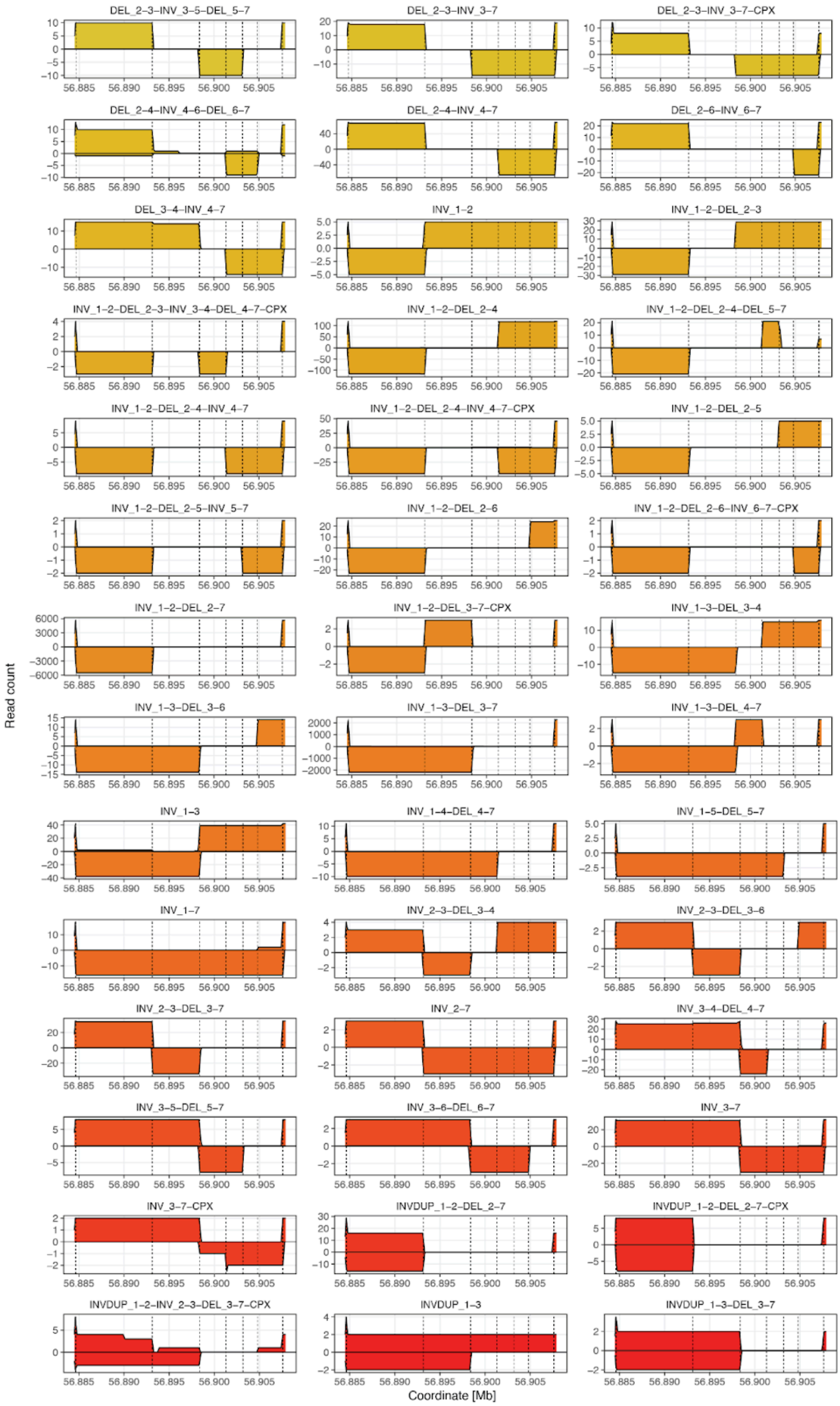


**Supplementary Figure 11. Regulatory architectures identified from scrambling OTX2-loxp7 cells.** Following recombination, cell sorting, PCR amplification, and sequencing, reads supporting specific genotypes were grouped (**Methods**), resulting in 134

unique synthetic architectures. (**a**) Read coverage (y-axis) across the scrambled region (x-axis) for architectures without inversion events. (**b**) As (a), but for architectures with inversion events. Negative read counts indicate coverage on the other strand. Dashed lines: positions of the 7 loxPsym sites. Architectures are named based on the type of rearrangement (DEL: deletion, INV: inversion, and INVDUP: inverted duplication) and the loxPsym sites (1 to 7) involved in the scrambling reaction. Those labeled as complex (CPX) denote architectures where the super-enhancer domains have been rearranged and are no longer in their original order.

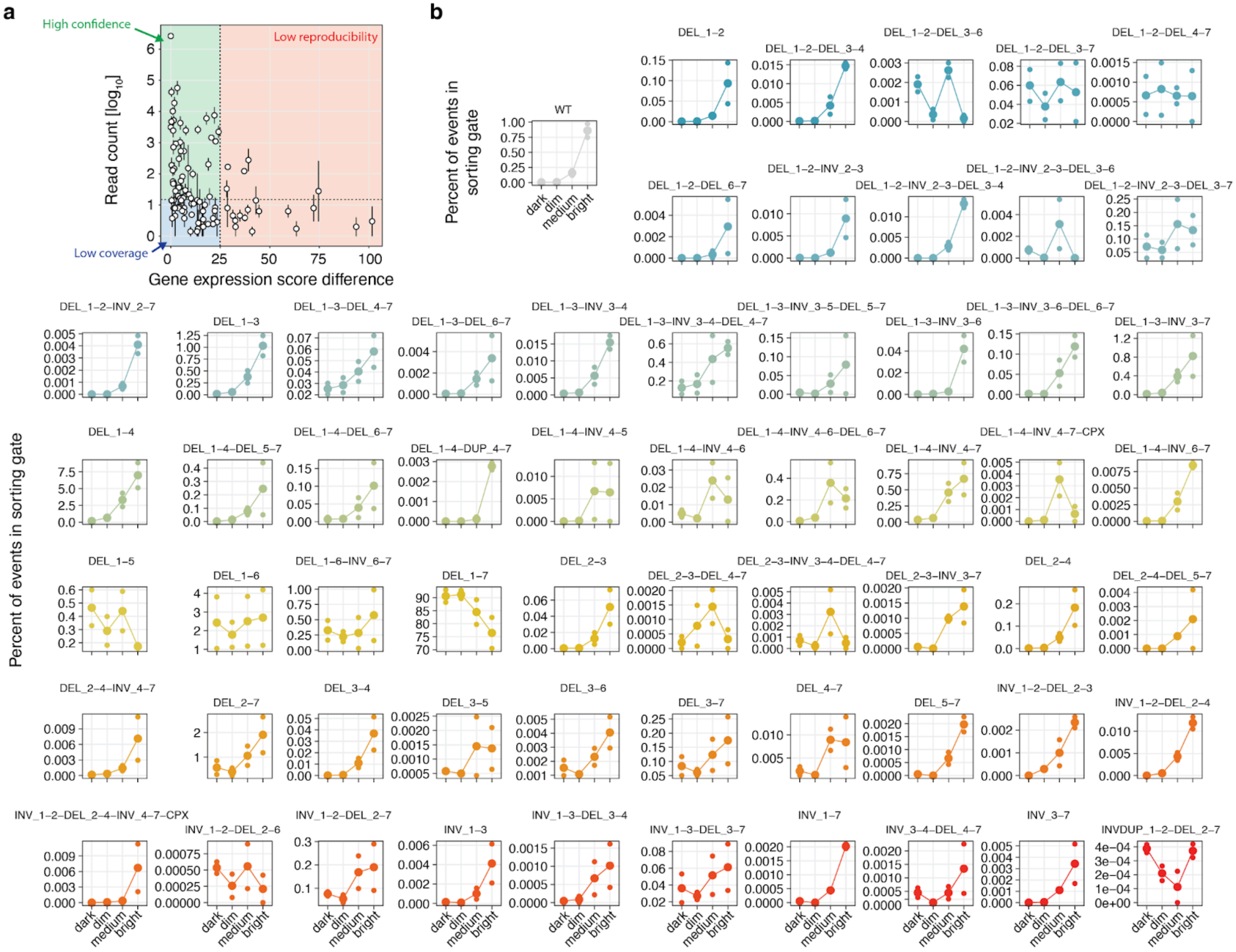
a
High confidence
Low reproducibility
Low coverage
Read count [log10]
Gene expression score difference
b
Percent of events in sorting gate
WT
dark
dim
medium
bright
DEL_1–2
DEL_1–2–DEL_3–4
DEL_1–2–DEL_3–6
DEL_1–2–DEL_3–7
DEL_1–2–DEL_4–7
DEL_1–2–DEL_6–7
DEL_1–2–INV_2–3
DEL_1–2–INV_2–3–DEL_3–4
DEL_1–2–INV_2–3–DEL_3–6
DEL_1–2–INV_2–3–DEL_3–7
DEL_1–2–INV_2–7
DEL_1–3
DEL_1–3–DEL_4–7
DEL_1–3–DEL_6–7
DEL_1–3–INV_3–4
DEL_1–3–INV_3–4–DEL_4–7
DEL_1–3–INV_3–5–DEL_5–7
DEL_1–3–INV_3–6
DEL_1–3–INV_3–6–DEL_6–7
DEL_1–3–INV_3–7
DEL_1–4
DEL_1–4–DEL_5–7
DEL_1–4–DEL_6–7
DEL_1–4–DUP_4–7
DEL_1–4–INV_4–5
DEL_1–4–INV_4–6
DEL_1–4–INV_4–6–DEL_6–7
DEL_1–4–INV_4–7
DEL_1–4–INV_4–7–CPX
DEL_1–4–INV_6–7
DEL_1–5
DEL_1–6
DEL_1–6–INV_6–7
DEL_1–7
DEL_2–3
DEL_2–3–DEL_4–7
DEL_2–3–INV_3–4–DEL_4–7
DEL_2–3–INV_3–7
DEL_2–4
DEL_2–4–DEL_5–7
DEL_2–4–INV_4–7
DEL_2–7
DEL_3–4
DEL_3–5
DEL_3–6
DEL_3–7
DEL_4–7
DEL_5–7
INV_1–2–DEL_2–3
INV_1–2–DEL_2–4
INV_1–2–DEL_2–4–INV_4–7–CPX
INV_1–2–DEL_2–6
INV_1–2–DEL_2–7
INV_1–3
INV_1–3–DEL_3–4
INV_1–3–DEL_3–7
INV_1–7
INV_3–4–DEL_4–7
INV_3–7
INVDUP_1–2–DEL_2–7

132

**Supplementary Figure 12. Impact of synthetic architectures identified from the scrambling of a heterogeneous OTX2-loxp7 cell population on *OTX2* expression. (a)** Difference in gene expression scores between replicates (x-axis) contrasted against the number of reads (y-axis; log10 scale) associated with the 100 identified architectures observed in both replicates (markers) following the scrambling of a heterogeneous OTX2-loxp7 cell population. Red: large differences between replicates (difference at least 25, "low reproducibility"); blue: low coverage (fewer than 15 reads); green: high coverage and high reproducibility ("high confidence"; **Methods**). Points, whiskers: replicate means and ranges. **(b)** Percentage of reads (y-axis), associated with the 61 high-confidence architectures (panels) within sorting gates (x-axis; gate definitions in **Supplementary Fig. 10)**. Architectures are named based on the type of rearrangement (DEL: deletion, INV: inversion, and INVDUP: inverted duplication) and the loxPsym sites (1 to 7) involved in the scrambling reaction. Architectures labelled as complex (CPX) refer to those in which the super-enhancer domains have been rearranged and are no longer in their original order.

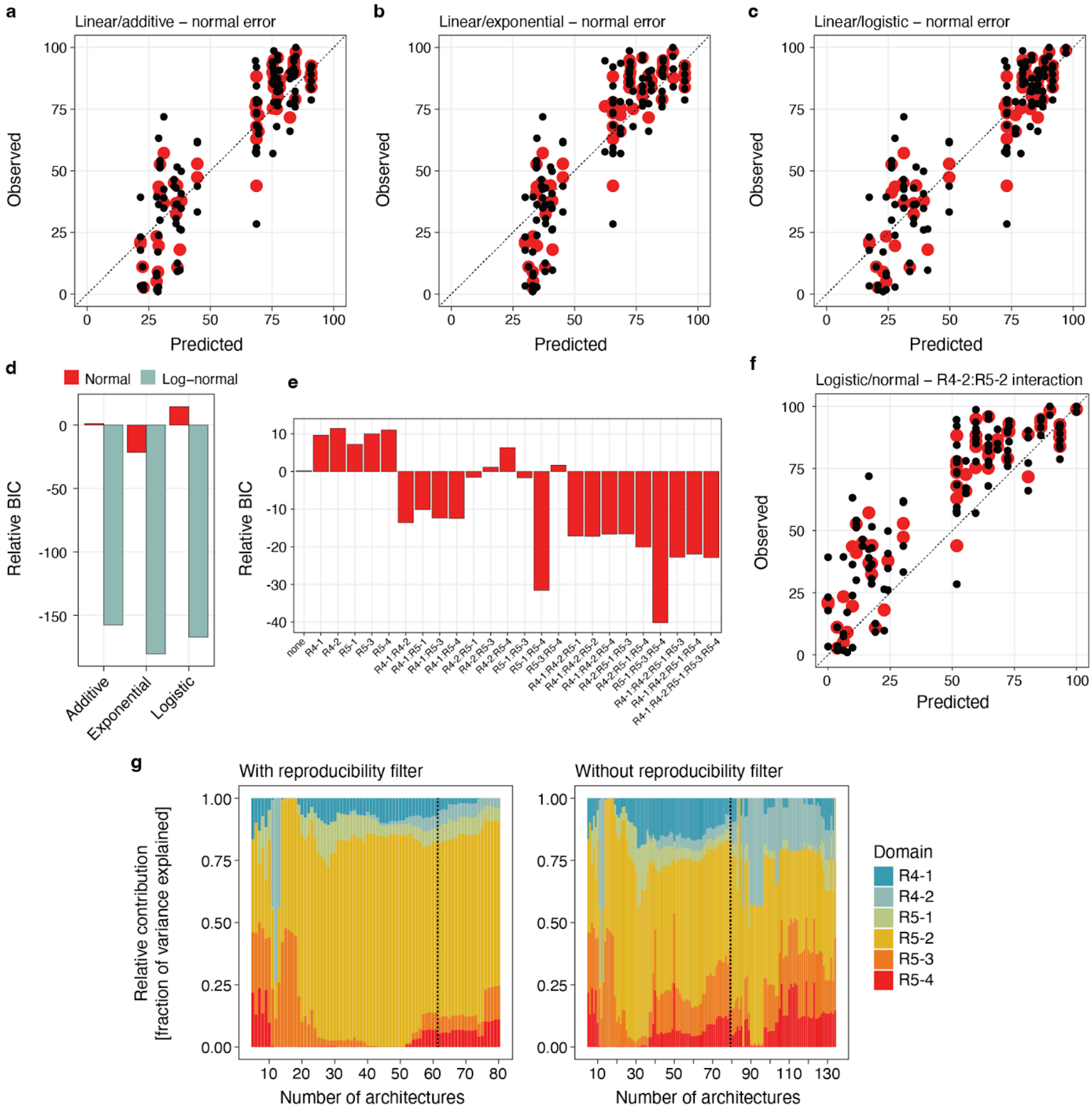


**Supplementary Figure 13. Modelling *OTX2* super-enhancer activity. (a)** Model fit (x-axis) and observed (y-axis) gene expression value of the 61 high-confidence architectures from scrambled OTX2-loxp7 cells (markers) using the linear/additive model with normal error. Black points: individual replicates; red points: replicate averages. **(b)** As (a), but for an exponential model. **(c)** As (a) but for a logistic model. **(d)** Bayesian information criterion relative to an additive model with normal errors (y-axis) of different alternative models (x-axis) and error models (colors). Red: normal errors; teal: log-normal errors. **(e)** As (d), but for models that allow interactions of different variable sets (x-axis) with the R5-2 domain. **(f)** As (a), but for the best-performing model, which considers interactions between the R4-2 and R5-2 domains. **(g)** Relative enhancer domain contributions (y-axis) computed from logistic-normal models applied to sets of architectures of increasing size, ranked by decreasing coverage (x-axis). Left panel: reproducibility filter applied; right panel: no reproducibility filter (**Methods**). Dotted vertical line: the last architecture set before reaching the coverage threshold used to define high-confidence architectures (Supplementary Fig. 12a).

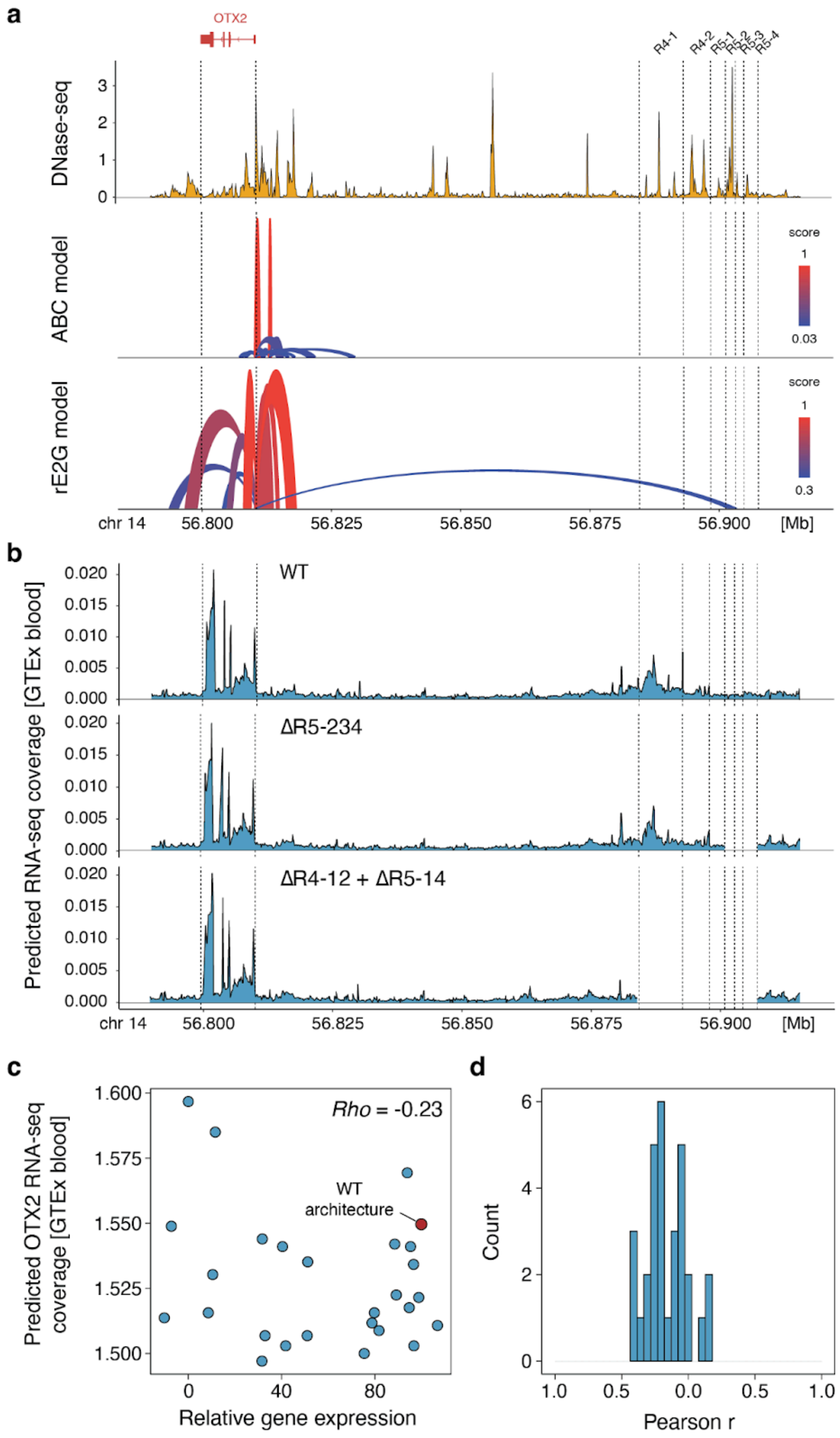


**Supplementary Figure 14. Predictions of enhancer interactions and effects of deletion variants on *OTX2* expression. a**) The *OTX2* locus accessibility (top panel; y-axis) for the gene (red box) and its scrambled regulatory domains (marked regions), complemented with Activity-by-Contact (ABC) model predictions (middle panel), and ENCODE-rE2G model predictions (bottom panel). Arches: predicted interactions; arch heights, colour: model score.

**b**) Predicted RNAseq coverage from the Borzoi model using the GTEX blood output (y-axis) along the OTX2 locus (x-axis) for wild-type (WT) configuration (top panel); and scrambled configurations ΔR5-234D (middle panel), and ΔR4-12+ΔR5-14 (bottom panel). **c)** Predicted RNA-seq coverage summed across the *OTX2* gene (y-axis) contrasted against measured gene expression score (x-axis) for scrambled regulatory configurations of simple deletions only (markers). Red marker: wild type architecture. Rho: Spearman correlation coefficient. **d**) all 31 cell types.

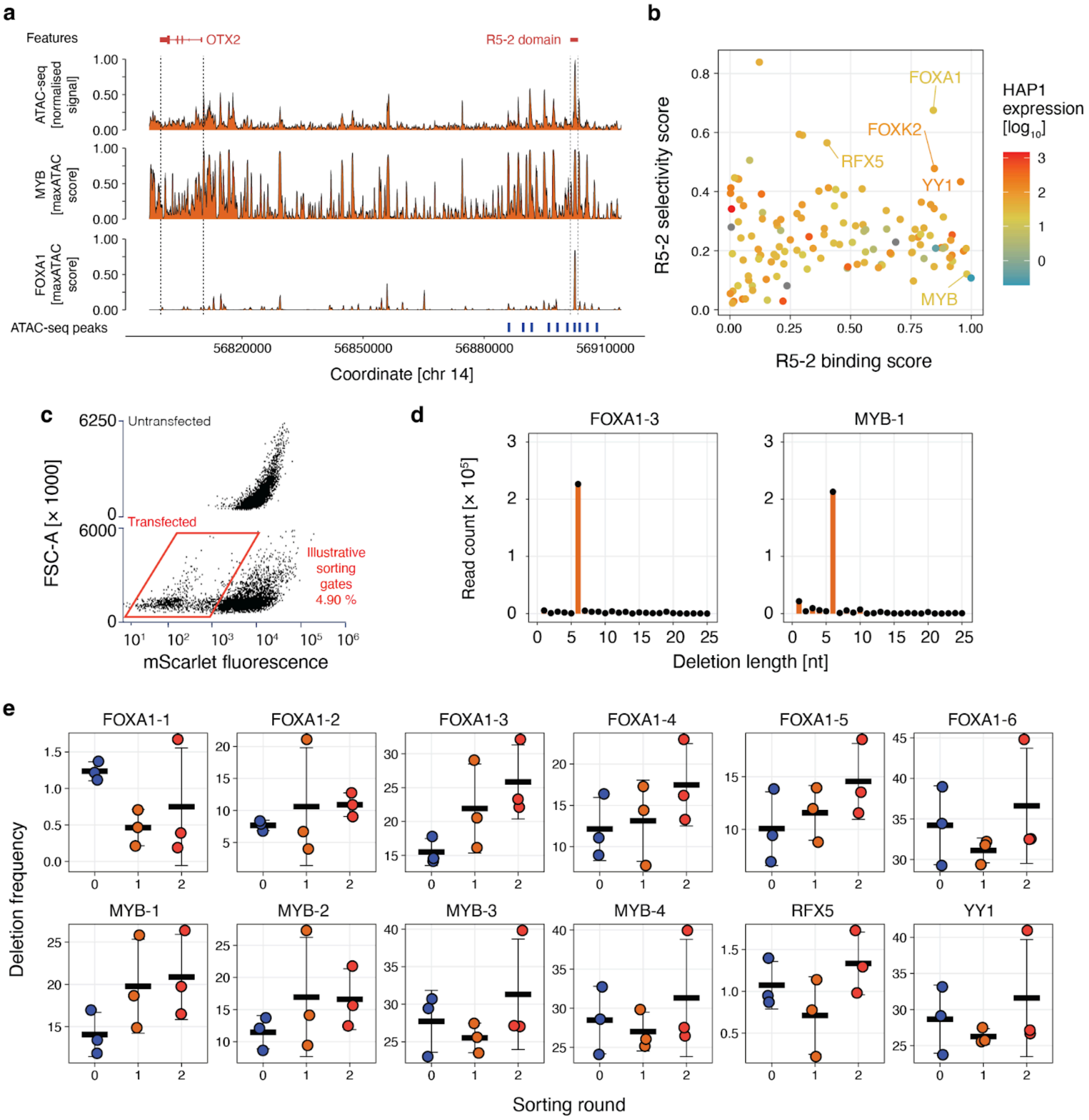


**Supplementary Figure 15. Targeting deletions at distal transcription factor binding sites. (a)** Genomic tracks displaying normalised ATAC-seq signals (y-axis; top panel) at the *OTX2* locus (x-axis) alongside maxATAC prediction scores for the FOXA1 (middle panel) and MYB (bottom panel) transcription factors (TFs). Red boxes: *OTX2* gene and the R5-2 domain of the *OTX2* super-enhancer. **(b)** R5-2 domain binding score (x-axis) and selectivity score (y-axis) for 135 human TFs expressed in HAP1 cells (markers). Highlights: FOXA1, FOXK2, YY1, and RFX5 as selective TFs, with MYB characterised as a more ubiquitous TF. **(c)** After two rounds of transfection with 12 epegRNAs targeting deletions at predicted binding sites of these TFs, flow cytometry identified a distinct population of transfected (bottom panel) but not untransfected (top panel) HAP1 cells (markers) with reduced

*OTX2-mScarlet* expression (x-axis) independent of cell size (forward scatter, y-axis). FACS was used to isolate approximately 5% of cells exhibiting low *OTX2-mScarlet* expression levels (red gate). Two rounds of sorting were conducted (**Methods**). **(d)** Intended deletions were detected within R5-2 amplicons generated from transfected cells. Number of reads (y-axis) with different deletion lengths (x-axis) highlight a large frequency of intended 6 nt deletions at targeted FOXA1-3 and MYB binding sites across all amplicons analyzed. **(e)** In addition to intended deletions, longer deletions (resulting from the concurrent prime editing activity of the 12 epegRNAs) overlapping with the targeted TF binding sites were identified. Frequency of deletion (y-axis) of the 12 binding sites (panels) for each round of sorting (x-axis, with round 0 representing no sorting). Lines and whiskers: means and standard deviations. Markers: individual replicates.

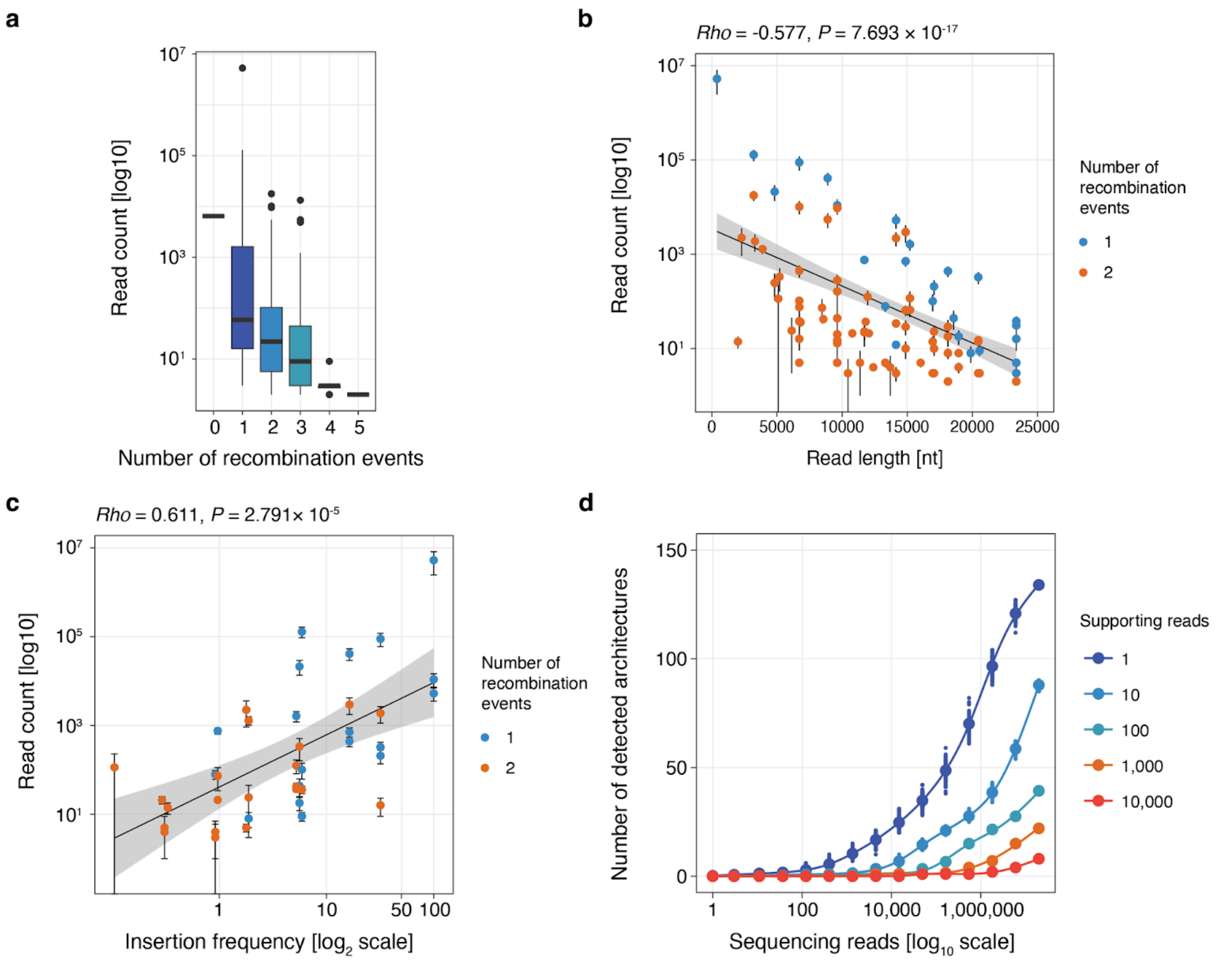


**Supplementary Figure 16. Scope and limitations of enhancer scramble. (a)** Number of reads (log10-scale, y-axis) supporting architectures with a number of recombination events needed to generate it (x-axis) for the 134 synthetic architectures obtained from the scrambling of a heterogeneous OTX2-loxp7 cell population. Box plots indicate medians and interquartile ranges. **(b)** Read count (log10-scale, y-axis) contrasted to read length (x-axis) for architectures (marker) generated from either one (blue) or two (orange) recombination events. Rho: Spearman's correlation coefficient. **(c)** Read count (y-axis, log10 scale) contrasted to estimated insertion frequency of all involved loxPsym sequences (x-axis) at sites involved in the generation of deletion structural variants from the scrambling of OTX2-loxp7 cells. Colors: number of breakpoints. Whiskers: range across replicates. Linear regression lines with confidence intervals are shown by the line and shaded area. *Rho:* Pearson's correlation. **(d)** Average number of detected architectures from resampling reads at a fixed coverage 100 times (y-axis) as a function of sequencing depth (x-axis, log10 scale).

### Supplementary Tables

**Supplementary Table 1.** Sequence of oligonucleotides and primers used in this study.

| Application | Sequence |
|---|---|
| mScarlet insertion<br>Forward primer | GACCTTCCCTCCCTTCCTTCAC |
| mScarlet insertion<br>Reverse primer | CTTCCTACTTTGGGGGCATGGA |
| *OTX2* targeting crRNA for T2A-mScarlet tagging | ctacaggtcttcacaaaacc |
| DNA donor used to tag *OTX2* with T2A-mScarlet tagging | tttgtccatttcatgttgctggtttgtaggcccctctaaggcccttcgtt<br>tttccttctatgcctctcggaactttgatcagatgagtctgagcatca<br>tcccatctaactcttttaaccaatgcctggctaaaactggaatgtc<br>cagcccagtatatttaaaaatcacccacaaaaagaggttctaca<br>ggtcttcacaaaACTTTATACAGCTCGTCCATCCC<br>GCCTGTGCTATGTCTACCTTCACTCCGCTCG<br>TACTGCTCAACTACAGTATAATCTTCGTTGTG<br>AGAGGTAATGTCCAATTTGCGATCCACATTAT<br>AGGCCCCGGGCATCTGCACAGGCTTCTTGG<br>CTTTGTAGGTTGTTTTAAAGTCGGCCAGGTA<br>TCTCCCACCGTCCTTTAGGCGCAGTGCCAT<br>CTTTATGTCACCCTTAAGCACGCCATCTTCA<br>GGGTACAGCCTCTCCGTAGAGGCCTCCCAC<br>CCCATAGTTTTCTTCTGCATAACTGGCCCGT<br>CGGGCGGGAAATTGGTTCCCCTCAGTTTAA<br>CTTTGTAAATTAACGTGCCGTCTTCGAGTGA<br>AGTATCCTGAGTCACGGTGACTGCCCCACC<br>ATCCTCGAAGTTCATGACCCGTTCCCACTTG<br>AAGCCTTCAGGGAAGCTTTGCTTATAGTAAT<br>CGGGGATATCAGCAGGATGCTTGGTGAACG<br>CACGGGATCCGTACATAAATTGTGGGCTCAA<br>GATGTCCCATGAAAATGGAAGGGGGCCTCC<br>TTTTGTGACCTTGAGTTTAGCTGTCTGGGTG<br>CCTTCATATGGTCGTCCCTCGCCCTCTCCCT<br>CGATTTCGAACTCGTGTCCGTTCATGGAGCC<br>TTCCATGTGCACTTTAAACCTCATAAACTCCT<br>TAATTACAGCCTCGCCCTTACTCACCATGGG<br>GCCGGGGTTCTCCTCGACGTCTCCGCAAGT<br>GAGCAGACTGCCTCTCCCCTCACCGCCGGA<br>GCCTCCAacctggaatttccacgaggatgtctgatctttataa<br>tccaagcagtcagcattgaagttaagcttccaggaggcagtttg<br>gtccttataatccaagcaatcagtggttgagttaaaacccaagctt<br>gaagctccatatccctgggtggaaagagaagctggggactgat<br>tgagatggctggtgactgcattggtacccat |
| Amplification of loxPsym insertion site chr14: 56,884,634. | TGGTTGTTGGAGGTGGGTGGGG |
| Amplification of loxPsym insertion site chr14: 56,884,634. | GGATGGCGTATGAGCGGGATGC |
| Amplification of loxPsym insertion site chr14: 56,898,376. | TGCCTCCTCCTCTCATGAAACCT |
| Amplification of loxPsym insertion site chr14: 56,898,376. | GCAAAACGGCTCAGACAACCCCA |

| | |
|---|---|
| Amplification of loxPsym insertion site chr14: 56,907,608. | AGACAATGTCCCTGCCCTCAAG |
| Amplification of loxPsym insertion site chr14: 56,907,608. | ACCAGCATTGCTTGGAAGTGTT |
| Amplification of loxPsym insertion site chr14: 56,840,403. | TCTCCTTCCACTCTGATTGCTCT |
| Amplification of loxPsym insertion site chr14: 56,840,403. | AGCATAGAAAGTGGCTGGAGCT |
| Amplification of loxPsym insertion site chr14: 56,851,953. | TAGTCGCAGTTACTTGGGAGGC |
| Amplification of loxPsym insertion site chr14: 56,851,953. | TTAGCACAAGGGCCAGAAATGC |
| Amplification of loxPsym insertion site chr14: 56,864,718. | AGTACTCACTTGGCAACCAGCT |
| Amplification of loxPsym insertion site chr14: 56,864,718. | CTCCTGGCAAGGGAAGGAAAGA |
| Amplification of loxPsym insertion site chr14: 56,884,634. | TGGTTGTTGGAGGTGGGTGGGG |
| Cas9-enrichment of the 20 kb enhancer cluster. 5’ probe 1. | gttgaccatgttgaaccagg |
| Cas9-enrichment of the 20 kb enhancer cluster. 3’ probe 1. | tggtccaagtaaacaaacgg |
| Cas9-enrichment of the 20 kb enhancer cluster. 3’ probe 2 / Cas9-enrichment of the 70 kb region with 6 loxPsym sites. 3’ probe 2 | gttccataccaagcaagcag |
| Cas9-enrichment of the 70 kb region with 6 loxPsym sites. 5’ probe 1. | gcagaaacacgaagtaacat |
| Cas9-enrichment of the 70 kb region with 6 loxPsym sites. 5’ probe 2. | agggtgcccattatagtggt |
| Cas9-enrichment of the 70 kb region with 6 loxPsym sites. 3’ probe 2. | cactcctcaaatgcactacc |
| Amplification of the OTX2 super-enhancer domain<br>Forward primer | TCTCAACATCAAAGAGCACCCCT |
| Amplification of the OTX2 super-enhancer domain<br>Reverse primer - index 1 | ggtgctgAAGAAAGTTGTCGGTGTCTTTGTGacc agcattgcttggaagtgtt |
| Amplification of the OTX2 super-enhancer domain<br>Reverse primer - index 2 | ggtgctgTCGATTCCGTTTGTAGTCGTCTGTacc agcattgcttggaagtgtt |
| Amplification of the OTX2 super-enhancer domain<br>Reverse primer - index 3 | gtgctgTTCGGATTCTATCGTGTTTCCCTAaccag cattgcttggaagtgtt |
| Amplification of the OTX2 super-enhancer domain<br>Reverse primer - index 4 | ggtgctgCTTGTCCAGGGTTTGTGTAACCTTacc agcattgcttggaagtgtt |
| Amplification of the OTX2 R5-2 domain<br>Forward primer - with illumina adapter | acactctttccctacacgacgctcttccgatctAGTCCCACC CTTAATCTCTTTGA |
| Amplification of the OTX2 R5-2 domain<br>Reverse primer - with illumina adapter | gtgactggagttcagacgtctgctcttccgatctTCATTCATG TTTGATTCAGCACA |

212

**Supplementary Table 2.** Sequence of epegRNAs used in this study**.**

| Protospacer | Extension | Application |
|---|---|---|
| GCTAGAGTAGGGCAGCTACA | cagcaatgactcctacctgtggatccctgtATAACTTCGTATAATGTACATTATACGAAGTTATagctgccctactc | Insertion of loxPsym<br>Insertion site chr14: 56,840,403 |
| GAATTTAGAGCCCTACGAGG | gctgacagagacaggcggtaggatccacctATAACTTCGTATAATGTACATTATACGAAGTTATcgtagggctctaa | Insertion of loxPsym<br>Insertion site chr14: 56,851,953 |
| GATTGGTCCAGAATGCCCAT | atctctcaggctacgccaatcctacctatgATAACTTCGTATAATGTACATTATACGAAGTTATggcattctggacc | Insertion of loxPsym<br>Insertion site chr14: 56,864,718 |
| GGACTACCATCTATCTGTGT | tgaaaggacgtttgtgatcccagccccacaATAACTTCGTATAATGTACATTATACGAAGTTATcagatagatggta | Insertion of loxPsym<br>Insertion site chr14: 56,884,634 |
| ACTCTTGAGTCAAGTCACCT | tcagaattgtgaagacgttatgaacccaggATAACTTCGTATAATGTACATTATACGAAGTTATtgacttgactcaa | Insertion of loxPsym<br>Insertion site chr14: 56,893,129 |
| GACATCAAATGTACCCCAGT | acttcacaaaattaagggtttctgccaactATAACTTCGTATAATGTACATTATACGAAGTTATggggtacatttga | Insertion of loxPsym<br>Insertion site chr14: 56,898,376 |
| TGCTGCGATTGGTGAGCTAG | gaagtcctcctcattgcacccccacccctaATAACTTCGTATAATGTACATTATACGAAGTTATgctcaccaatcgc | Insertion of loxPsym<br>Insertion site chr14: 56,901,292 |
| TTCATGGTGTGTCATGCCGG | ttactttatagttgtaatgatcccccgATAACTTCGTATAATGTACATTATACGAAGTTATgcatgacacacca | Insertion of loxPsym<br>Insertion site chr14: 56,903,183 |
| AAAGAAGGAATGCAGGTTAG | ttactatctgttatccccaccccacccctaATAACTTCGTATAATGTACATTATACGAAGTTATacctgcattcctt | Insertion of loxPsym<br>Insertion site chr14: 56,904,792 |
| GAGGGCATTCTAAGAGTTAG | ttcaacttcaagaaatcctctattccctaATAACTTCGTATAATGTACATTATACGAAGTTATactcttagaatgc | Insertion of loxPsym<br>Insertion site chr14: 56,907,608 |
| TGGTGTGGGCCACACATATA | ACTTTACAAAAGATTGAGGAAAGGATTGGACAAACTTTCCTTATATGTGTGGCCCACA | Deletion of FOXA1-1 site<br>T[CTTTAT]T > TT |
| TTGCCCCATCTCCTTTAGTA | AACATTATGCAAGGGCATAGGAATAAGGCCTTACTAAAGGAGATGGGG | Deletion of FOXA1-2 site<br>G[TAAATA]C > GC |
| TCTTTTCATTCTCTGAGCAG | AGTCGCCACTTATATGCACTTCAATCCGCTGCTCAGAGAATGAAAA | Deletion of FOXA1-3 site<br>A[TTTTTA]C > AC |
| CTCTGCAGCTTCGGCCCAGT | TCAGAGAATGAATCTGGACTTAAGCTGTTTACCAACTGGGCCGAAGCTG | Deletion of FOXA1-4 site<br>A[AAGATG]T > AT |

| | | |
|---|---|---|
| GTTTTGCCGCTCTGCAGCTT | GTCTGGACTTAAGCCCAACTGG GCCGAAGCTGCAGAGCGG | Deletion of FOXA1-5 site C[TGTTTA]C > CC |
| GCTTAGAAGAGTTTCAGGGT | AGTTGAGTCCCACCCTGAAACT CTTCTA | Deletion of FOXA1-6 site T[TATTAA]C > TC |
| GCTTAGAAGAGTTTCAGGGT | TTTACAATGCGAGCAGCAGTTG AGTTATTAACCCACCCTGAAACT CTTCTA | Deletion of YY1 site G[CCATTTT]C > GC |
| AGATCTTTTGTAAAGTGCTA | TGAGGCCTCTATGAGTCACTTTA CAAAAGA | Deletion of RFX5 site T[TTCCATAG]C > TC |
| TAATGTTACAGGCTATTGTT | ATATAAGTGGACTGGCGCTGGC CTAACAATAGCCTGTAA | Deletion of MYB-1 site T[CAGTCG]C > TC |
| GTTTTGCCGCTCTGCAGCTT | CTGGACTTAAGCTGTTTACGGC CGAAGCTGCAGAGCGG | Deletion of MYB-2 site C[CAACTG]G > CG |
| ACTGGGCCGAAGCTGCAGAG | TAAAAGCTTGTTCCTGCTTGGG GCTTTCTTCAGTTTTGCCGCTC TGCAGCTTCGGCCCA | Deletion of MYB-3 site G[CAACTG]C > GC |
| GCTTAGAAGAGTTTCAGGGT | AATGCGAGCAGCCATTTTAGTTA TTAACCCACCCTGAAACTCTTCT A | Deletion of MYB-3 site T[CAGTTG]A > TA |

214 cr772 scaffold: gtttaagagctaagctggaaacagcatagcaagtttaaataaggctagtccgttatcaactcgaaagagtggcaccgagtcggtg.
